# Gut-relevant short-chain fatty acids modulate host-pathogen dynamics of uropathogenic *Escherichia coli* at the colonic epithelial interface

**DOI:** 10.64898/2026.01.19.700340

**Authors:** Nicholas Yuen, Ramon Garcia Maset, Wendy Wang, Manas Kubal, Zekun Chen, Victoria Chu, Ian J. White, Jinhui Gao, Francesca Torelli, Aaron Crowther, Laia Pasquina-Lemonche, Jennifer L. Rohn

**Affiliations:** Center for Kidney and Bladder Health, Division of Medicine, University College London, London WC1E 6BT UK; Institute of Biomedical Engineering, Department of Engineering Science, University of Oxford, Oxford, UK; Laboratory for Molecular Cell Biology, University College London, London WC1E 6BT, UK; Institute of Biomedical Engineering, Department of Engineering Science, University of Oxford, Oxford, UK; Nuffield Department of Orthopaedics, Rheumatology and Musculoskeletal Science, University of Oxford, Oxford, UK; School of Biosciences, University of Sheffield, Sheffield, UK

## Abstract

Urinary tract infection (UTI) ranks among the most prevalent bacterial infections worldwide, affecting over 400 million people each year. Uropathogenic *Escherichia coli* (UPEC), the main aetiological cause of UTI, colonises the intestinal tract, which is thought to serve as a distal reservoir for gut-UTI recurrence. Despite this, the precise role of the gut in UTI recurrence is still not fully defined. Recent research investigating the gut-UTI axis has revealed that reduced abundance of gut commensals producing short-chain fatty acids (SCFAs, namely acetate, butyrate and propionate) is associated with recurrent and chronic UTI. We therefore aimed to investigate the impact of these gut commensal-derived metabolites on a diverse panel of UPEC strains, including well-studied prototypical strains (UTI89, CFT073), a non-pathogenic isolate *E. coli* K-12, and various clinical UTI isolates (from the urine of both symptomatic and asymptomatic individuals). We observed that SCFAs modulate bacterial growth kinetics in a concentration- and pH-dependent manner, by prolonging the lag phase without affecting final carrying capacity *in vitro*. These metabolites further suppressed bacterial swimming motility and biased the orientation of *fimS*, the phase variable switch for T1 fimbriae, under acidic conditions. In a human polarized, mucus-secreting intestinal infection model, SCFA treatment during UPEC challenge altered bacterial localization patterns, favouring planktonic over mucosal-associated populations, and preserved epithelial barrier function. Together, these *in vitro* findings demonstrate that SCFAs modulate key UPEC colonization-associated phenotypes and influence host-pathogen dynamics at the colonic epithelial interface. These results provide mechanistic insights into how depletion of SCFA-producing gut commensals may alter the intestinal reservoir environment *in vitro* and warrants further investigation into the role of gut-derived SCFAs in rUTI susceptibility.

## Introduction

Urinary tract infection (UTI) is among the most common bacterial infections globally, affecting over 400 million people worldwide each year (Yang et al., 2022). Although UTI is more frequent in women, with 50% experiencing at least one infection in their lifetime, men and children are also affected. These infections often recur, with up to 25% of women experiencing another UTI within six months (Aydin et al., 2015), thought to be caused in part by treatment failure due to the host/pathogen interaction (Garcia-Maset et al., 2025). As a result, UTI creates a substantial economic burden, with an estimated annual cost of US$5 billion in the United States alone (Dudzik et al., 2025). The rising prevalence of antimicrobial resistance (AMR) further worsens the clinical situation, with studies estimating that ∼260,000 UTI deaths were associated with AMR in 2019 globally (Li et al., 2022). Collectively, these data highlight UTIs as a substantial clinical challenge, and further research is needed to elucidate the mechanistic underpinnings of recurrence. However, like many diseases affecting predominantly or solely women, UTI remains an understudied area (Head et al., 2020).

Both Gram-negative and Gram-positive bacteria can cause UTI. Among these, uropathogenic *E. coli* (UPEC), a pathotype of extra-intestinal *E. coli,* accounts for approximately 80-90% of community-acquired and 40-50% of hospital-acquired infections (Flores-Mireles et al., 2015; Kucheria et al., 2005). One major factor thought to play a role in recurrent infections is the intestinal reservoir, in which UPEC can shelter between bladder episodes. Current evidence suggests that the gut acts as the primary reservoir for many uropathogens identified in UTI patients. Despite this, whether the gut reservoir acts as a passive bystander, facilitator or agitator is not fully understood (Worby et al., 2022a). Whole genome sequencing of *E. coli* from healthy controls (faecal) and UTI patients (faecal and urinary samples) indicates that these isolates are closely related and differ mainly in their accessory genomes, with limited variation in the core genome as determined by analyses of genomic single nucleotide polymorphisms (Nielsen et al., 2017). More recent studies using 16S rRNA and metagenomic sequencing of faecal samples have reported that even a 1% abundance of intestinal uropathogens is a significant UTI risk factor, with further analysis showing strain-level similarity between gut and urine isolates from the same patient (Magruder et al., 2019). Longitudinal studies using clonal tracking have demonstrated that an intestinal bloom of UPEC often precedes and facilitates repeated transfer of uropathogens between the intestinal reservoir and the urinary tract within the same individuals, highlighting the causal relationship between gut expansion of UPEC and subsequent UTI (Thänert et al., 2022, 2019). Despite evidence that intestinal abundance correlates with susceptibility to cystitis, conflicting findings in the literature suggest that additional factors influence this relationship (Iqbal et al., 2024).

Taken together, the current gut-UTI paradigm indicates that uropathogens residing in the gut share genomic homology with those found in the bladder, likely serving as a source of infection in recurrent UTI. However, other reservoirs, such as the bladder and vagina, may also play important roles in recurrence (Hunstad and Justice, 2010; Lewis and Gilbert, 2020). Thus, recurrence is likely multifactorial, with the gut serving as an important contributor. As the gut represents a primary reservoir for UPEC, the characteristics of this reservoir become critical to understanding recurrence risk. Emerging evidence suggests that it is not simply the presence of UPEC in the gut, but the state of the surrounding microbial community and metabolic environment, that determines whether strains are poised for recurrent infection. Thus, the intestinal environment, particularly during dysbiosis, may significantly influence rUTI susceptibility (Meštrović et al., 2021).

One hypotheses points to the modulation of bacterial physiology and virulence. UPEC possesses multiple virulence factors that help to facilitate infection, often acquired through chromosomal mutations, mobile genetic elements and/or acquisition of pathogenicity islands encoding multiple advantageous genes (Bunduki et al., 2021; Zagaglia et al., 2022). One key category includes an arsenal of adhesins, which enable UPEC to attach and colonize multiple sites along the urinary tract (Flores and Rohn, 2025).

The mechanisms by which UPEC colonize and persist within the intestinal niche is an area of active research. Recent evidence suggests that mucosal association in the gastrointestinal tract is predominantly mediated by type 1 pili (Azimzadeh et al., 2025), whereas *Ucl* and *Yeh* pili were found to bind exclusively to faecal content in the colonic lumen. Deletion of *fimH*, the tip adhesin of the type 1 pilus, was further found to result in upregulation of *fliC*-mediated swimming motility in both human and mouse inner mucus layer ex-vivo. Azimzadeh *et al*. further proposed that this compensatory mechanism may drive UPEC to breach the inner mucus layer to access the epithelium. Thus, both type 1 adhesins and motility have been described as key colonisation traits which may serve to drive recurrence and persistence within the gut niche.

Given the reported importance of these colonization traits for UPEC persistence in the gut, an important next question is how features of the intestinal microenvironment, particularly those altered during dysbiosis, shape the regulation of these virulence programs. Emerging evidence indicates that women with recurrent UTI (rUTI) frequently exhibit gut microbiota disruption, including a marked reduction in short-chain fatty acid (SCFA)–producing taxa such as *Faecalibacterium spp*., *Akkermansia muciniphila*, *Blautia*, *Romboutsia spp*., and *Eubacterium halii* (Worby et al., 2022b). Consistent with this, higher abundances of *Faecalibacterium* and *Romboutsia* species have been associated with reduced *Enterobacteriaceae* bacteriuria and lower UTI risk in kidney transplant patients (Magruder et al., 2019). SCFAs are metabolic by-products of anaerobic fermentation carried out by these commensal taxa. These beneficial metabolites play key roles in host physiology by serving as energy sources, regulating gene expression, reinforcing gut barrier integrity, and promoting anti-inflammatory pathways (Mirzaei et al., 2022).

SCFAs also directly influence bacterial physiology, affecting growth and transcriptional programs in *E. coli* (Kadry et al., 2023; Zhang et al., 2020) as well as in other pathogens such as Klebsiella, Salmonella and *Staphylococcus aureus* (Fletcher et al., 2025; Lawhon et al., 2002; Sorbara et al., 2019). Notably, Young *et al*. reported that intestinal *E. coli* from women with rUTI exhibit a transcriptional shift toward aerobic metabolism, including altered expression of key regulators ArcA and FNR, suggesting metabolic adaptation, likely driven by depletion of SCFA-producing taxa (Young et al., 2024). This altered transcriptional program may facilitate increased likelihood of successful bladder infections. Together, these findings indicate that rUTI patients possess *E. coli* with distinct transcriptomic states that, when associated with certain microbiome contexts, may predispose them to enhanced UTI recurrence. Despite the emerging connection between dysbiosis, SCFA depletion, and altered *E. coli* physiology, how SCFAs modulate the colonization traits of UPEC within the gut remains poorly understood. Defining these SCFA-driven effects on UPEC host-pathogen interactions should provide valuable insight into the therapeutic role of these beneficial metabolites.

## Methods

**Table 1:**
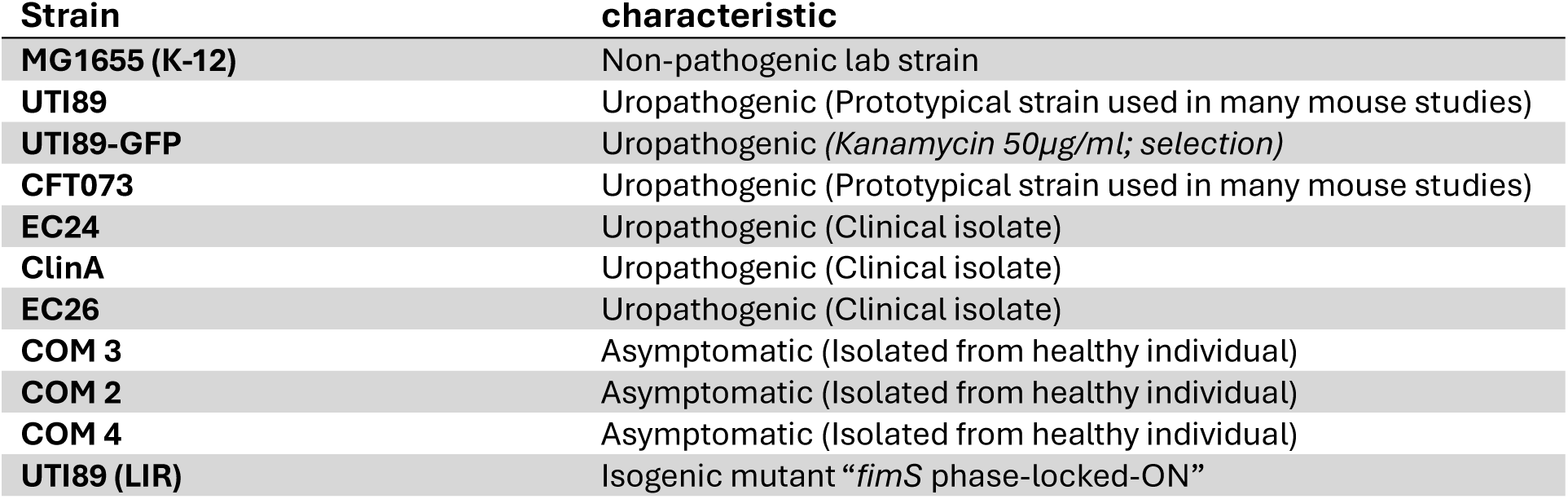
Bacterial Strain List.

### Media

Bacteria were routinely grown in 2.5% Lysogeny Broth (LB, Sigma) media and plated on LB with 1.5% Lennox agar (Sigma). The SCFA mix was prepared by making a 1 M total solution (ddH20) of sodium acetate (Merck), sodium butyrate (Insight Biotechnology), and sodium propionate (Fisher Scientific) at a molar ratio of 60:20:20, respectively, to reflect typical SCFA levels and composition (Den Besten et al., 2013). pH was adjusted via titration with 1 M hydrochloric acid (Fisher Chemical) or sodium hydroxide (Reagecon) and was measured with a Mettler Toledo SevenEasy pH meter, calibrated as instructed by the manufacturer. UTI89-GFP was cultured in LB supplemented with kanamycin (50 mg/mL).

### Swimming Motility Assay

Single colonies of bacterial strains were grown overnight on LB agar plates. On the day, soft agar containing 1% tryptone, 0.5% NaCl, and 0.25% LB agar was autoclaved (Merino and Tomás, 2016). Afterwards, sterile 1 M SCFA or 1 M NaCl (Thermo Fisher) stock solution was added to the cooled agar solution immediately before pouring, resulting in a final concentration of 120 mM. Plates were left to dry for 1 hr in sterile conditions after which a single colony of each bacterial strain was used to stab-inoculate the agar (Chen et al., 2026), followed by incubation of the plates at 37°C for 18 hours. Motility was quantified by measuring the diameter of the growth of each strain, or the longest distance from end to end when not perfectly circular. Three biological experiments were performed with three technical replicates per experiment.

### Growth Curves

Bacterial cultures were made by inoculating single colonies from fresh agar plates into 5 ml of LB broth and grown in an orbital shaking incubator overnight at 37°C at 200 RPM. The following day, bacterial overnight cultures were adjusted to an OD_600_ of 0.1 and further diluted 1:100 in LB media supplemented with either SCFA mixture, individual SCFAs or NaCl at final concentrations of 120 mM, 60 mM and 20 mM. NaCl was used as the untreated control following previously published work on this method (Zhang et al., 2020). Two hundred µl of inoculum was added to each well in a 96-well plate in triplicate, and OD_600_ measurements were taken every 30 minutes over 17.5 h with a Tecan Spark® 10M microtiter plate reader at 37°C with orbital shaking (180 rpm, 3 mm amplitude) every 5 minutes. Three biological experiments were performed with three technical replicates per experiment.

### *FimS* Inversion PCR-Restriction Fragment Length Polymorphism (RLFP) assay

Bacterial strains were cultured under fimH phase-ON inducing conditions. A single colony of *E. coli* UTI89 or CFT073 was inoculated in 5 mL of LB with and without SCFA mixture (120 mM final concentration) at pH 6.2 and 7.2 and cultured overnight in static conditions at 37°C. After 24 h overnight cultures were subcultured 1:1000 in their respective media and grown for another 24-hour in static conditions at 37 °C, as previously described (Greene et al., 2015). Following incubation, cultures were standardised to an OD_600_ of 1 in. PCR amplification of FimS was performed with 1 µl of overnight culture (OD_600_ = 1) with Phusion High Fidelity polymerase (ThermoFisher) with previously validated primers (Forward primer: AGTAATGCTGCTCGTTTTGC; Reverse primer: GACAGAGCCGACAGAACAAC) (Chen et al., 2014). Briefly, each PCR sample contained 0.5µM of each primer, 0.02 U/µL of Phusion polymerase, 200µM dNTPs and 1x Phusion HF Buffer in PCR-grade water. The following PCR program was used: 98°C 30 s; 30 cycles of (98°C 10 s; 60°C 30 s; 72°C 18 s); 72°C 600 s; 4°C. To validate the assay, the UTI89 wild type (WT) and the fimS phase-locked-ON UTI89 mutant, which has point mutations in its left inverted repeat (Kostakioti et al., 2012) and cannot be reoriented into the OFF phase (both kindly provided by Scott Hultgren’s laboratory), were used as controls (Greene et al., 2015). Both WT and mutant strains were grown under Phase-OFF conditions cultured on LB agar plates, followed by fimS PCR of resuspended colonies. The PCR product was digested with 10 units of SnaBI (Jena Bioscience) in a 50ul reaction for 37 °C for 2 hours and inactivated at 80°C for 20 minutes. 10 µL of digested product with BlueJuice Gel Loading Buffer (Invitrogen) was run on a 2% agarose gel with SYBR Safe DNA Gel Stain (Invitrogen) in TAE buffer, alongside a 0.5 µL of GeneRuler 100 bp DNA Ladder (Invitrogen). Gel electrophoresis was performed at 120 V for 90 minutes, and bands were visualised using the Invitrogen iBright Imaging System (Invitrogen). ImageJ analysis was performed to determine the relative brightness of bands, and %ON and %OFF was calculated as (ON/OFF)/(ON+OFF) *100. Experiments were performed in 5 biological replicates.

### Establishment of a <u>Co</u>lonic <u>M</u>ucosal/microtissue In-vitro Model for <u>M</u>icrobial–Intestinal <u>C</u>rosstalk (Co-MIMIC)

Caco2 (human colonic enterocyte-like cells) and HT29-MTX-E12 (human Goblet-like cells, hyper mucus secreting sub-clone) were purchased from the European Collection of Cell Culture (ECACC) and received at passage number 1 and 51 (Lot Number HT29-MTX-E12: 22B018) respectively. Cells were routinely grown in T150 flasks (Corning) in 24 ml of pre-warmed supplemented DMEM (Thermo Fisher) (High glucose, L-glutamine, Pyruvate negative, supplemented with 10% heat-inactivated FBS (Thermo Fisher), 1% MEM Non-essential amino acids (Thermo Fisher), 1% Pen/Strep (Thermo Fisher) with 3 media changes weekly until they reached a 70-80% confluency, then passaged with 0.25% trypsin 1 mM EDTA (Thermo Fisher) dissociation at 37^°^C for 4 mins after 1x PBS wash.

Resuspended cells were spun at 300 g for 5 minutes and either sub-cultured at a 1×10^5^ density or seeded for the intestinal epithelial model. Passages between 20-27 and 60-70 were used for Caco2 and HT29-MTX-E12 cells respectively when seeding onto 12 mm diameter 0.4 µm pore size polycarbonate Snapwell inserts (Corning, REF3407) in co-culture at a 3:1 ratio (7.5×10^4^:2.5×10^4^) to reflect a higher proportion of goblet cells in the colonic region (Tóth et al., 2017). 500 µL and 1.5 mL of media were added to the apical and basolateral compartments respectively and changed every 2-3 days for 14 days until mature.

### CellTracker Staining

For validation of Co-MIMIC morphology, HT29-MTX-E12 cells were stained with CellTracker green dye (Invitrogen) prior to seeding in the Co-MIMIC system. A freshly thawed CellTracker green vial was left to equilibrate at room temperature for 5-10 minutes, before dilution at a final concentration of 25 µM in serum-free pre-warmed DMEM media. Cells were resuspended in 1 ml CellTracker green solution and incubated for 45 minutes at 37^°^C, before being spun, resuspended and seeded in the Co-MIMIC model as usual. This was left to incubate for 45 mins and cells were resuspended. Post-incubation, cells were centrifuged and resuspended in DMEM media and used for co-culture seeding in Snapwells (Corning). Membranes were mounted with ProLong Glass Antifade mounting solution (Invitrogen) onto glass slides and imaged with a confocal microscope (Leica SP8). Images were processed with the Leica Application Suite (LASX) and ImageJ (Fiji) Software.

### Alcian Blue Staining

An Alcian Blue staining kit (ABCAM PLC) was used validate expression of acidic mucins as previously described (Xavier et al., 2019). Briefly, 4% methanol-free formaldehyde (Invitrogen)-fixed model co-cultures were excised from Snapwells using a scalpel and submerged in ultrapure water for 5 minutes. Membranes were subsequently submerged in acetic acid solution for 3 minutes, followed by Alcian Blue stain (pH 2.5) for 30 minutes at RT. Next, membranes were washed twice with distilled water and counter-stained with nuclear fast red solution for 5 minutes. After two more washes with distilled water, samples were dehydrated through graded ethanol: 70% (1 minute), 95% (1 minute), and 100% (2 minutes). Finally, membranes were mounted onto glass slides using ProLong Glass Antifade mounting media (Invitrogen) and imaged with a Leica inverted microscope DMi1 (Leica Microsystems).

### Cell Count Quantification for MOI estimation

To determine cell density for multiplicity of infection (MOI) calculations, mature Co-MIMIC samples were fixed and stained as in the below “Immunofluorescence” section, followed by confocal imaging. Confocal Z-stacks (imaged at 40x magnification) of the 4′,6-diamidino-2-phenylindole stained (DAPI; nuclei) channel were used to determined cell counts using the TrackMate plugin workflow in FIJI (Tinevez et al., 2017). To quantify nuclei counts, the following parameters were applied: an estimated spot diameter of 8 μm, Laplacian of Gaussian (LOG) filter for spot detection, and a linking distance and gap-closing distance set to 1 μm. The Z-stack dimension was used to represent sample depth. Spots corresponding to nuclei were identified based on the specified diameter and linked through the stack using these parameters (**Supplementary Figure S1**). The total number of unique tracks generated corresponded to the cell count for each sample (four randomly selected regions were imaged across three independent experiments) in the given area and extrapolated to estimate the number of cells across a 1.12 cm^2^ polycarbonate membrane.

### Bacterial infection and Quantification in Co-MIMIC

Prior to infections, UTI89-GFP was cultured at 37°C for 48 h in static conditions to induce type 1 fimbriae expression as described previously (Schilling et al., 2001). In DMEM without sodium bicarbonate, bacterial cultures were adjusted to an OD₆₀₀ of 0.1 and diluted 1/100, of which 500 µl were inoculated into the apical compartment to correspond to an MOI of 0.2 (total cell number in the Co-MIMIC model is ∼2.5 × 10^6^). 1.5 mL of fresh DMEM supplemented with sodium bicarbonate was added to the basolateral compartment, and the cultures were incubated at 37°C in 5% CO2. 14 hours post infection (hpi), the apical compartment (containing planktonic bacteria) was carefully collected, ensuring minimal disruption to the mucus layer. To remove any residual planktonic bacteria, the apical compartment was gently washed once with an equal volume of sterile PBS. To quantify adhered/ intracellular bacteria, the remaining epithelial tissue was lysed using 1% Triton X-100 diluted in PBS for 15 minutes at 37°C. Both were serially diluted (log₁₀) and plated on LB agar in 10 µL spots for colony-forming unit (CFU) counts the next day. For fixation, Co-MIMIC samples were treated with 4% methanol-free formaldehyde (Invitrogen) for 15 minutes at room temperature, after which samples were washed and stored in PBS for downstream processing. Three to four biological experiments were performed with three technical replicates per experiment.

### Immunofluorescence Staining and Microscopy

Following fixation with either 4% methanol-free formaldehyde or methacarn fixative (60% methanol, 30% Chloroform, 10% glacial acetic acid) for preservation of mucus, membranes were excised from Snapwells using a scalpel and blocked with 3% BSA in PBS (Merck) for 30 minutes at room temperature. The following staining was performed in 3% BSA for 90 minutes on gentle rocking prior to permeabilization to detect mucins: mouse anti-MUC2 monoclonal antibody conjugated to AlexaFluor 488 (Santa Cruz Biotechnology sc-515032 AF488, 2 µg/mL). For staining of intracellular proteins, membranes were permeabilized with 0.2% Triton X-100 (Sigma-Aldrich) for 20 minutes, before blocking in 3% BSA for 30 minutes. The following were then added: mouse anti-ZO-1 monoclonal antibody conjugated to AlexaFluor 647 (Life Technologies MA3-39100-A647, 5 µg/ml), Phalloidin conjugated to Alexa Fluor 555 (Invitrogen) and 4′,6-diamidino-2-phenylindole (DAPI, Invitrogen, 2 μg/ml). Membranes were mounted with ProLong Glass Antifade mounting solution (Invitrogen) onto glass slides and imaged with a confocal microscope (Leica SP8) at 40X and 60X magnification. Images were processed with the Leica Application Suite (LASX) and ImageJ (Fiji) Software.

### Scanning Electron Microscopy (SEM)

Membranes for SEM processing were fixed in methacarn fixative (60% methanol, 30% Chloroform, 10% glacial acetic acid) for 30 minutes at room temperature, prior to processing and imaging at the Electron Microscopy facility at the Laboratory for Molecular Cell Biology at UCL. Fixed membranes were incubated with 1% osmium tetroxide (TAAB Laboratories)/1.5% potassium ferricyanide (Sigma-Aldrich) for 1 hour at 4°C, followed by three washes with 0.1 M CAC buffer. Subsequently, they were incubated in 1% tannic acid (TAAB Laboratories) in 0.05 M CAC buffer in the dark at room temperature for 40 min, followed by two washes with 0.05 M CAC buffer and one wash with distilled and deionized water (ddH2O). Membranes were then dehydrated in graded ethanol (Sigma-Aldrich).

Dehydrated microtissues were dried with a Leica EM CPD300 critical point dryer. SEM images were acquired using a Zeiss Gemini 300 with a working distance of 8 mm, 1.5 kV accelerating voltage using a secondary electron (SE2) detector and processed with Zeiss Atlas 5 software.

### Trans-Epithelial Electrical Resistance (TEER)

TEER was measured to assess barrier integrity. Briefly, an STX4 electrode (World Precision Instruments) was sterilised in ethanol then rinsed in ultrapure water for 5 minutes and conditioned in DMEM for 5 minutes before TEER measurement. In preparation, fresh pre-warmed DMEM was added to all Snapwells with 500 µl in the apical compartment and 4 ml in the basolateral compartment. All samples, including a blank control (Snapwell containing no cells), were equilibrated at 37°C with 5% CO_2_. The STX4 electrode was connected to a EVOM3 Epithelial Volt/Ohm Meter (World Precision Instruments). A custom 3D-printed holder was used to ensure the electrode was placed correctly in the Snapwell insert (https://github.com/Degra97/STX2-Plus-Transwell-Holder). Corrected TEER was adjusted for the area of the culture according to the following formula: (TEER _actual_ – TEER _blank_) × 1.12 cm². Four biological experiments were performed with two technical replicates for infected samples and 7 biologicals with 1 technical for uninfected samples.

### FITC-Dextran Permeability Assay

Paracellular permeability was determined by the flux of fluorescein isothiocyanate (FITC)-labeled dextran MW 4000 (FD-4) (Sigma Aldrich; St. Louis, MO, USA) through the Co-MIMIC coculture after 14 days. The monolayer was washed with PBS and pre-equilibrated with Hank’s balanced salt solution (HBSS; Gibco) buffered to pH 7.4 at 37°C for 1 hour. Cells were carefully washed twice with PBS and 200 µL FD-4 solution (1 mg/mL in HBSS) was added to the apical side of the monolayers. Samples (100 µL) were collected from the basal chamber at 2,4,6,8 and 24 hours. 100 µL of fresh media was added at each sample collection to replace media taken. The fluorescent intensity of these basolateral aliquots was determined using the Tecan Spark microplate reader. Wavelengths of excitation and emission were 490 and 530 nm respectively. Three biological experiments were performed with two technical replicates each.

### Cell Cytotoxicity

The amount of released lactate dehydrogenase (LDH) in the apical compartment was measured as a proxy for cell death. A 2X LDH assay solution was prepared with 0.88 mM 2-p-iodophenyl-3-p-nitrophenyl-5-phenyl tetrazolium chloride (INT, Cayman Chemical 16073), 0.37 mM N-methylphenazonium methyl sulfate (PMS, MP Biomedicals 100955), 1.73 mM nicotinamide adenine dinucleotide (NAD, Sigma Aldrich 1245420001) and 57 mM L-lactic acid (Sigma Aldrich L7022-5G) in 200 mM Tris buffer solution pH 8.0, and aliquoted at -20°C until use. Briefly, apical supernatants were collected, centrifuged to remove cellular debris, and the supernatants were diluted 1:10 in PBS. For each sample, 50 µL of diluted supernatant was mixed with 50 µL of LDH reaction buffer in a 96 well plate and incubated for 30 minutes without light. The reaction was stopped by adding 1 M acetic acid and incubated for a further 15 min. Absorbance was measured at 490 nm and 680 nm with a Tecan Spark plate reader, and values were background-corrected by subtracting A680 from A490. Three biological experiments were performed with two technical replicates per experiment.

### Image Analysis and Quantification of ZO-1 Coverage

Maximum-intensity projections of the far-red channel, corresponding to anti-ZO-1 monoclonal antibody conjugated to AlexaFluor 647, were generated from confocal *z*-stacks acquired using a Nikon Ti2 confocal microscope with a 40× objective. Segmentation was performed using Ilastik (Berg et al., 2019). A supervised random forest-based machine-learning pixel classifier to generate segmentation masks of ZO1 staining was used. The model training was performed on a representative image with 1240×1240 pixels for each condition (i.e., one for SCFA treated and untreated). The trained model was then applied in batch to the rest of the images from each dataset. The resulting masks were binarized in FIJI, and the ZO1 coverage area was quantified as the proportion of positive pixels relative to the total image area. To quantify the mean length of the objects from the binarized mask, the AnalyzeSkeleton plugin (Arganda-Carreras *et al*., 2010) in Fiji was used and the branch length was reported. Six images were analysed per biological replicate (*n* = 3).

### Statistical Analysis and Schematics

Data were expressed as mean ± standard deviation, plotted, and analysed using GraphPad Prism version 10.5.0. Statistical significance between groups was assessed using analysis of variance (ANOVA). QURVE (Wirth *et al*., 2023) software version 1.1 was run in RStudio version 2024.09.1+394 and used to analyse time-resolved growth curve data. To obtain robust estimates of UPEC growth dynamics, we applied complementary parametric and non-parametric (for validation) analysis pipelines. Parametric analyses were performed using automatic model selection based on Akaike information criterion (AIC) model fitting to estimate key growth parameters (lag time, maximum growth rate, and carrying capacity). In parallel, non-parametric methods (e.g., spline fitting or area-under-the-curve comparisons) were applied to avoid assumptions regarding curve shape. All schematics were created with BioRender.com

## Results

### SCFAs modulate UPEC growth kinetics in a pH- and concentration-dependent manner

Recently, rUTI has been linked with alterations in gut microbiota composition, particularly of those taxa that secrete SCFAs. To investigate this, we evaluated the impact of the three predominant SCFAs – acetate, butyrate and, propionate – and their combination on the cystitis-associated UPEC strain UTI89. First, we assessed UTI89 growth dynamics exposed to SCFAs (single or in combination) ranging from 20 to 120 mM at either pH 6.2 or 7.2, conditions representative of the large intestinal environment spanning the proximal and distal colon (Den Besten et al., 2013; Yamamura et al., 2023). We included an equimolar NaCl control to account for osmotic balance as previously described (Zhang et al., 2020). Our results show that the addition of SCFAs buffered to a pH of 6.2 caused a dose-dependent growth inhibition, with maximal suppression at 120 mM and minimal suppression at 20 mM (**Fig. 1a; Supplementary Figure S2**), with a clear delay on the lag phase. Among the three SCFAs, propionate displayed the strongest inhibitory effect at pH 6.2 at the same concentration. Notably, propionate also reduced final carrying capacity at both pH 6.2 and pH 7.2 at concentrations of 60 mM and 120 mM (**Fig. 1a; Supplementary Figure S2**).

**Figure 1.**
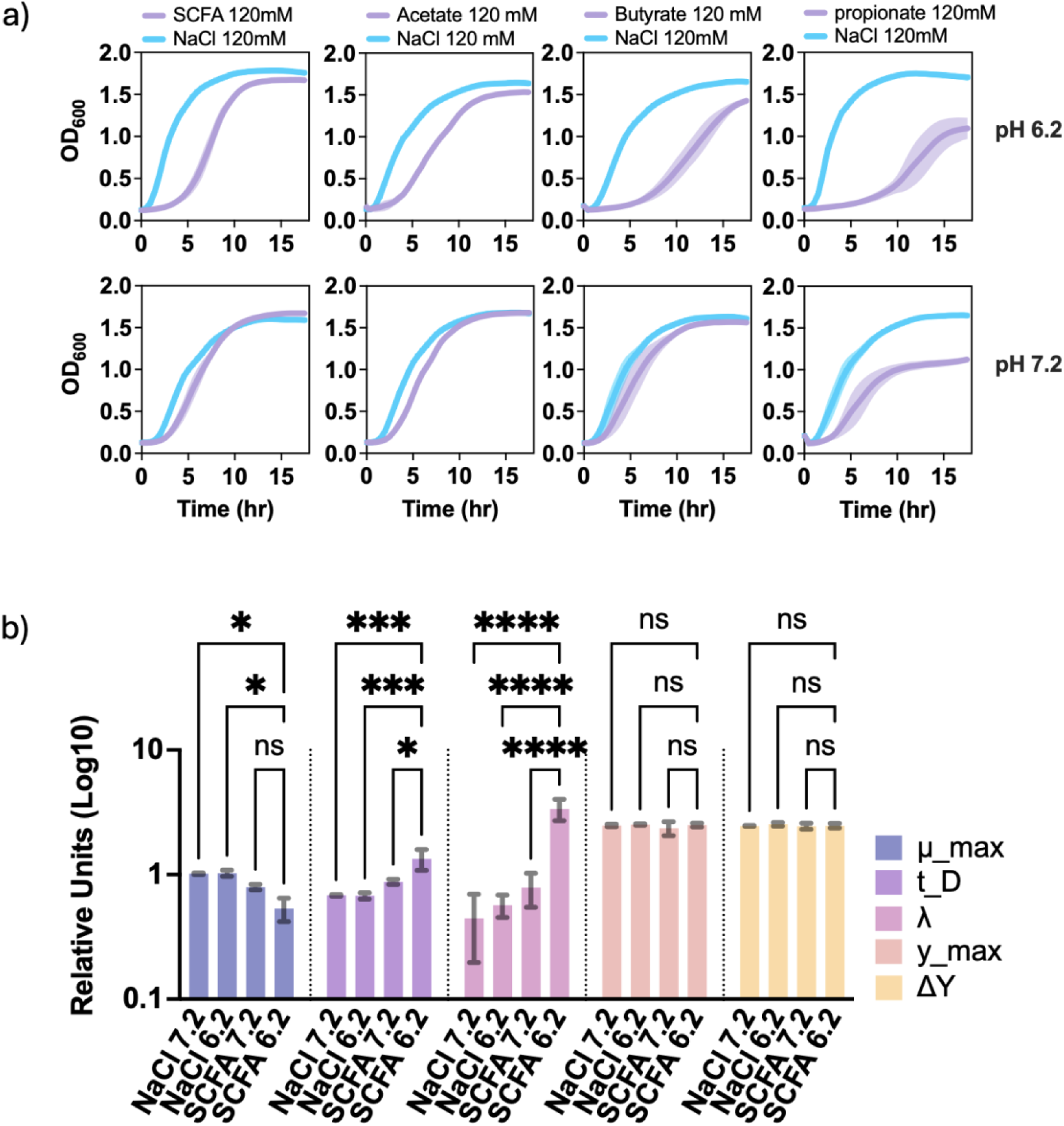
SCFAs modulate growth kinetics of uropathogenic *E. coli* strain UTI89 in a pH- dependent manner. **a**) Growth curves analysis of uropathogenic *E. coli* UTI89 grown in LB supplemented with SCFA mixture (Acetate: Butyrate: Propionate, 60:20:20 molar ratio), Acetate, Butyrate and Propionate at 120 mM and NaCl (at 120mM as positive control) at pH 6.2 and 7.2. Plots represent the mean and the error bars (shadow) shown as ± standard deviation of the mean. **b)** Quantitative (parametric fitting) comparison of UTI89 growth parameters under different pH and SCFA conditions. The following parameters are displayed in log10 relative units with error bars indicative of ± standard deviation from the mean: maximum specific growth rate (μ_max, blue), doubling time (t_D, purple), lag phase duration (λ, pink), maximum optical density (y_max, orange), and change in optical density (ΔY, yellow). An ordinary two-way ANOVA with Dunnett’s multiple comparisons test was performed to compare the different growth conditions within each parameter. Statistical significance is indicated above the bars (ns = not significant; *p<0.05; **p<0.01, ***p<0.001; ****p<0.0001). Data represent three biological and three technical replicates.

To further interpret our preliminary analysis, growth kinetic parameters were subsequently quantified when UTI89 was exposed to SCFA mixture at 120 mM (the maximal concentration tested) and the equivalent NaCl dose (**Fig. 1b**) For quantification, we used Qurve software to extract growth parameter data. Briefly, data were log-transformed and fitted against both parametric (**Fig. 1b**) and non-parametric models (**Fig. Supplementary S3, Supplementary Figure S4**) to derive kinetic parameters. We compared five key growth parameters: maximum growth rate (μ_max), doubling time (t_D), lag phase duration (λ), maximum cell density (y_max), and change in OD_600_ (ΔY) across four conditions tested (NaCl pH 7.2, NaCl pH 6.2, SCFA pH 7.2, and SCFA pH 6.2) in UTI89.

SCFA mix treatment significantly decreased μ max at pH 6.2 (P < 0.05) but not at pH 7.2 compared with NaCl controls. Conversely, μ max did not differ notably between the two NaCl conditions. Doubling time (t_D) was significant delayed by SCFA at both pH levels compared with NaCl, with the longest t_D observed under SCFA at pH 6.2. SCFAs also significantly increased the lag phase (λ) at pH 6.2 compared with all other conditions (p < 0.0001). However, maximum cell density (y_max) and overall change in OD_600_ (ΔY) were unaffected by either SCFA or pH. When analysing the data with non-parametric fitted models, similar patterns emerged with a significantly lengthened lag phase consistent with the parametric analysis (**Supplementary Figure S3**). We also observed that time to maximum growth rate t(μ_max) was higher in the SCFA pH 6.2 group, and area under the curve (AUC) was markedly reduced compared with all other conditions. Taken together, our findings indicate that SCFA exposure, especially at acidic pH, significantly prolonged the lag phase, while leaving the final cell biomass unchanged.

To assess whether this effect was conserved amongst other UPEC isolates, we tested the highest concentration of SCFA mixture on a panel of UPEC strains, including symptomatic and asymptomatic isolates from patient and healthy volunteer urine, respectively. All strains demonstrated a pH-dependent growth inhibition (**Fig 2a, b**), with significantly greater impairment observed at pH 6.2. Quantitative analysis of the AUC indicated that the acidified form of SCFAs was more effective as we observed a significant reduction in AUC, compared with SCFAs in neutral pH, in all strains tested. This reduction in AUC was observed to be attributed to a delayed lag phase, with eventual recovery to near-identical final densities. In contrast, growth inhibition was substantially reduced at pH 7.2 as determined by AUC, with significant reductions in only CFT073, MG1655, UTI89 and COM4 consistent with our initial observations in cystitis strain UTI89.

**Figure 2.**
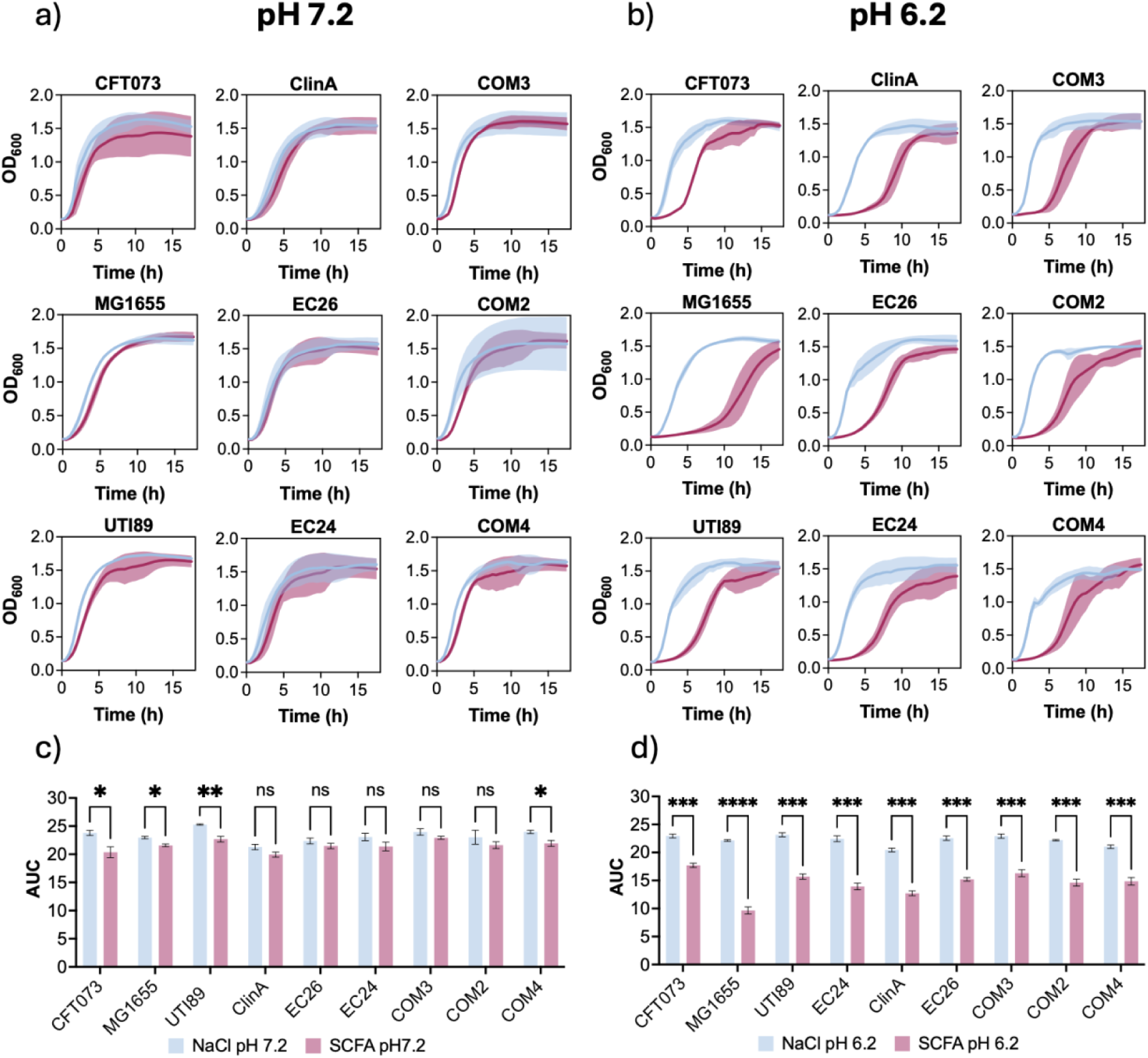
SCFA mixture impacts growth kinetics across multiple UPEC isolates including symptomatic and asymptomatic strains. Growth curves analysis examining the effects of SCFA supplementation in LB media across a panel of uropathogenic *E. coli* strains. Briefly, strains were grown in either 120 mM SCFA (Acetate: Butyrate: Propionate, 60:20:20) or 120 mM NaCl at a pH of 7.2 **a**) and 6.2 **b**). Area under the curve was quantified for all strains at pH 7.2 **c**) and 6.2 **d**). All data are representative of 3 biological and 3 technical replicates with error bars showing mean. Statistical analysis was determined by multiple unpaired t-test with statistical significance indicated above the bars: ns = not significant; *p<0.05; **p<0.01, ***p<0.001; ****p<0.0001.

### SCFAs attenuate swimming motility and fimS phase variation of UPEC

SCFAs have been shown to modulate virulence factors of other bacteria including *Salmonella spp*, *Klebsiella spp*, enterohemorrhagic *E. coli* (EHEC) and *Staphylococcus aureus* (Fletcher et al., 2025; Kadry et al., 2023; Lawhon et al., 2002). We tested whether SCFAs modulate key colonization phenotypes important for gut colonisation, such as those controlled by *fimH* and *fliC*. To assess the effect of SCFAs on swimming motility, we conducted swimming assays using soft agar plates supplemented with 120mM of either NaCl or the SCFA mixture at neutral pH (7.2). Quantitative analysis of motility zones revealed that SCFA induced a significant reduction in swimming diameters in all motile *E. coli* strains tested, including CFT073, MG1655, UTI89, EC24, ClinA, EC26, COM2, and COM4, compared with NaCl controls (**Fig. 3a, b**). No significant dispersion was observed in COM3, a non-motile strain. These findings demonstrate that SCFAs robustly reduced flagellar-driven swimming motility across a diverse panel of *E. coli* isolates, even under neutral pH conditions. These data suggest that SCFAs may directly interfere with colonisation-related behaviours independent of growth inhibition.

**Figure 3.**
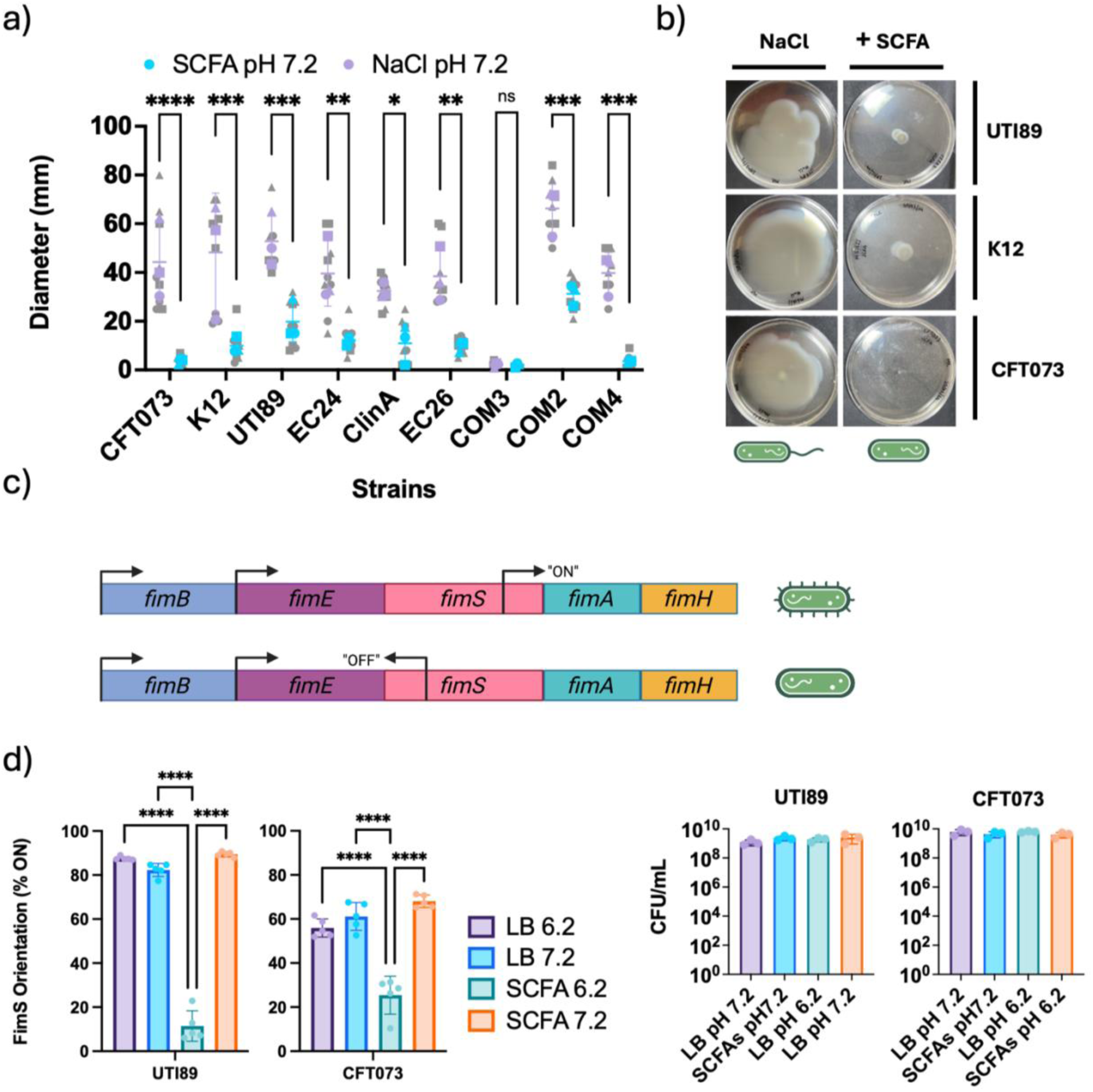
SCFAs inhibit flagella-dependent swimming motility and *fimS* phase variation in UPEC strains UTI89 and CFT073. **a)** Quantification of swimming diameters for a panel of *E. coli* strains grown on soft agar containing either NaCl (pH 7.2, blue) or physiologically relevant concentrations of short-chain fatty acids (SCFA, pH 7.2, purple). Individual data points represent 3 biological replicates (coloured) and 3 technical replicates (grey). Each strain plotted as mean ± SD. Statistical significance was assessed using an ordinary 2-way ANOVA with Holm-Šídák’s multiple comparison test. **b)** Representative images of swimming plates for three representative *E. coli* isolates (UTI89, CFT073, K12) after 16 hr incubation under NaCl or SCFA conditions at pH 7.2. c) Schematic of the *fim* gene cluster and *fimS* invertible element in the “OFF” (top) or “ON” (bottom) orientations, regulating downstream elements fimA and fimH in UPEC. d) Quantification of % *fimS* “ON” vs “OFF” orientation in two prototypical E. coli strains UTI89 and CFT073 grown in LB or SCFA-containing media at pH 7.2 (neutral) or 6.2 (acidic). Bar graphs show the %ON and %OFF orientation as mean ± SD (n=5). Statistical significance determined by two-way ANOVA; ns = not significant; *p<0.05; **p<0.01, ***p<0.001; ****p<0.0001. Created in BioRender. Yuen, C. (2026) https://BioRender.com/yvbgiqc

We further hypothesised that SCFAs play a role in modulating type 1 fimbriae. To test this, we analysed the orientation of the fimS invertible element, which controls the transcription of the downstream elements *fimA* and *fimH* (**Fig. 3c**), in two prototypical UPEC strains: UTI89 (cystitis) and CFT073 (pyelonephritis). Using a PCR-based phase variation assay, we quantified the relative abundance of *fimS* in the “ON” (pilus-expressing) and “OFF” (non-expressing) states under four different growth conditions: LB pH 7.2, LB pH 6.2, SCFA pH 7.2 (120 mM), and SCFA pH 6.2 (120 mM) (**Supplementary Fig S5**). For this assay an NaCl osmotic control was not included, as NaCl has been shown to directly regulate the fimS switch through effects on fimB/fimE (upstream promoter) transcription and fimS orientation, which would confound interpretation of fim-dependent outcomes. For UTI89 and CFT073, both LB conditions and the neutral SCFA yielded a higher fraction of cells with *fimS* in the “ON” state (**Fig. 3d**), relative to the acidified SCFA group (pH=6.2). To ensure similar viable input for PCR amplification, we quantified CFU/ml post OD_600_ normalisation (**Fig. 3d**). Acidification of the medium in the presence of the SCFA mixture resulted in a significant reduction of fimS “ON” orientation and a corresponding increase in the “OFF” state in both strains tested. These findings reveal that SCFAs in the protonated form robustly biases *fimS* switching to the “OFF” orientation, potentially repressing type 1 pilus expression across multiple UPEC backgrounds.

### Validation of Co-MIMIC

To investigate the effect of SCFAs on UPEC in the intestinal reservoir, we modified a published human gut-epithelial model (Anonye et al., 2019) to contain a proportion of goblet cells (HT29-MTX-E12; hyper mucus secreting subclone) to enterocyte ratio representative of the human colon, and to express an increased amount of mucus.

Firstly, we validated the epithelial organisation of co-MIMIC by performing morphological characterisation by optical and scanning electron microscopy (SEM) (**Figure 4**). Seven days after seeding, HT29-MTX-E12 goblet cells formed discrete clusters or “islands” surrounded by F-actin-rich Caco2 enterocytes (**Figure 4a, 4c S6a**), which was consistent with published models and ex-vivo histology (Walczak et al., 2015). We also observed mucus production by goblet cells, confirmed by MUC2 immunostaining and Alcian blue histochemistry (**Figure S6b**) consistent with previous reports (Xavier et al., 2019).

**Figure 4.**
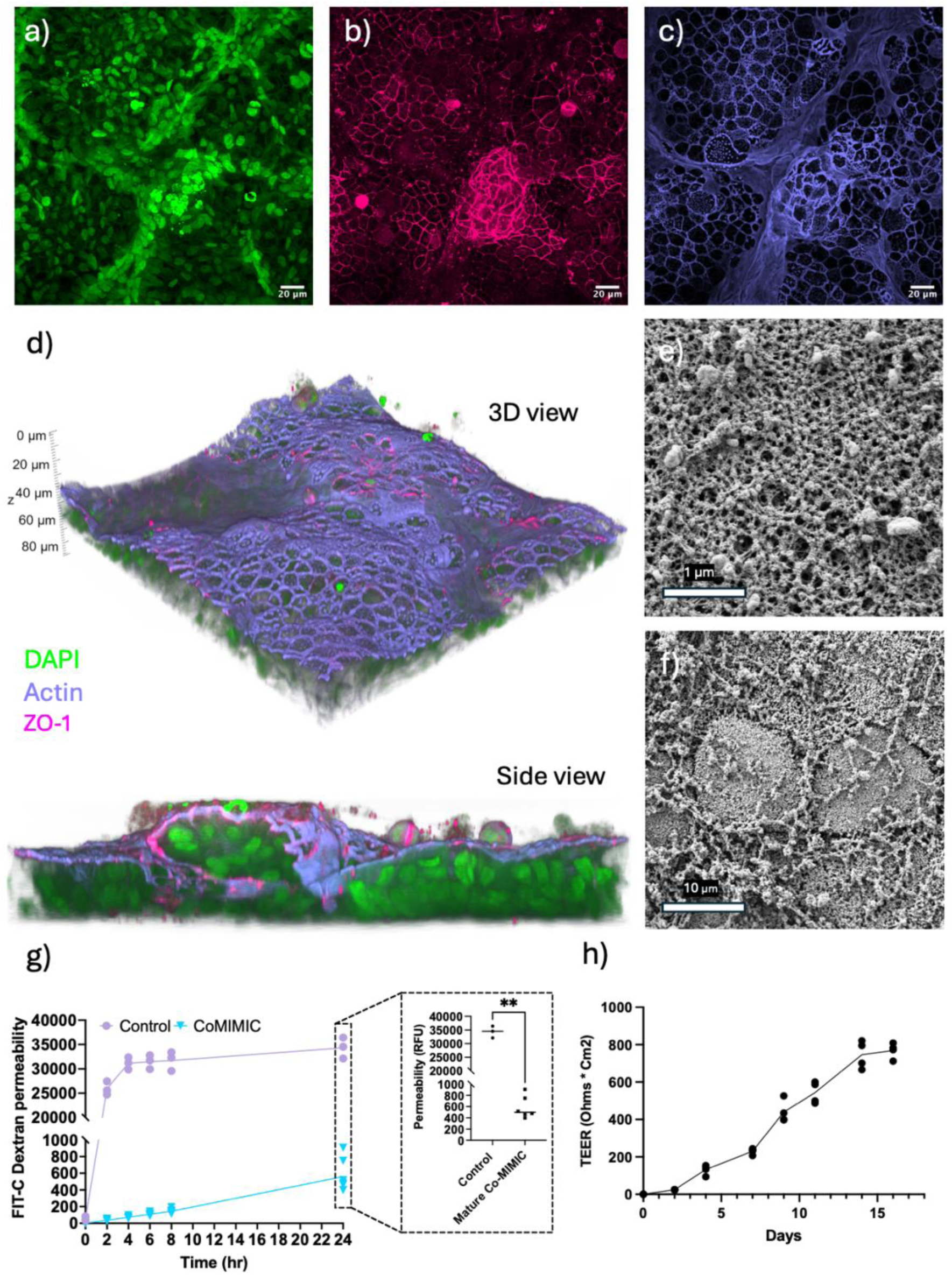
Morphological and molecular characterization of Co-MIMIC. Maximum-projection images of mature Co-MIMIC showing distribution of caco2 cells and HT29-MTX-E12 **a**) nuclei (green), localisation of **b**) ZO-1 tight junction protein (magenta) and **c**) actin (purple) **d**) 3D reconstruction of fully mature Co-MIMIC samples. **e**) SEM revealing microvilli structures on cell surface with **f**) epithelial organization. **g**) FIT-C dextran permeability assay comparing Co-MIMIC and no-cell control after 2 weeks. h) Trans-epithelial electrical resistance of Co-MIMIC maturation over time. All scale bars as indicated.

Furthermore, the tight junction protein ZO-1 localised continuously along cell borders of both Caco2 and HT29-MTX-E12 cells, suggesting that both cell types contribute to barrier integrity (**Figure 4b**). Three-dimensional reconstruction of confocal and SEM micrographs of the mature Co-MIMIC model revealed epithelial organisation and evidence of packed microvilli structures/brush borders, indicative of differentiation and polarization (**Figure 4e, f**). In addition, we determined enhanced barrier function via FIT-C permeability assay at endpoint (**Figure 4g**) as well as a gradual increase in TEER over time indicative of maturation (**Figure 4h**). Collectively, these findings confirmed that Co-MIMIC developed a physiologically organized intestinal epithelial interface.

### SCFAs mediate protection of the gut epithelium in response to UPEC challenge in a polarized and mucus-secreting human gut infection model

UPEC is widely considered to be a non-pathogenic coloniser of the intestinal niche, where it resides before traversing to cause infection in the bladder via the faecal-periurethral route (Worby et al., 2022). To investigate how SCFAs modulate UPEC colonisation in the intestinal niche, a 120 mM SCFA mixture was added during bacterial inoculation (t=0) onto our mature Co-MIMIC model for 14 hours under neutral or acidic conditions (**Fig. 5a**). SCFA supplementation significantly altered infection dynamics, resulting in more planktonic bacteria compared with that seen in the non-SCFA group independent of pH (**Fig. S7**). We observed no significant differences in adhered/intracellular bacterial burden across conditions (**Fig. S7**), suggesting that SCFAs promote a planktonic localisation but had a limited effect on epithelial association in this assay. When planktonic and adhered/intracellular fractions were expressed as % of total population, we observed that SCFA treatment increased the relative proportion of planktonic bacteria (∼50% SCFA vs 5% non-SCFA), while the non-SCFA group showed a predominantly mucosal-associated phenotype (**Fig 5b**).

**Figure 5.**
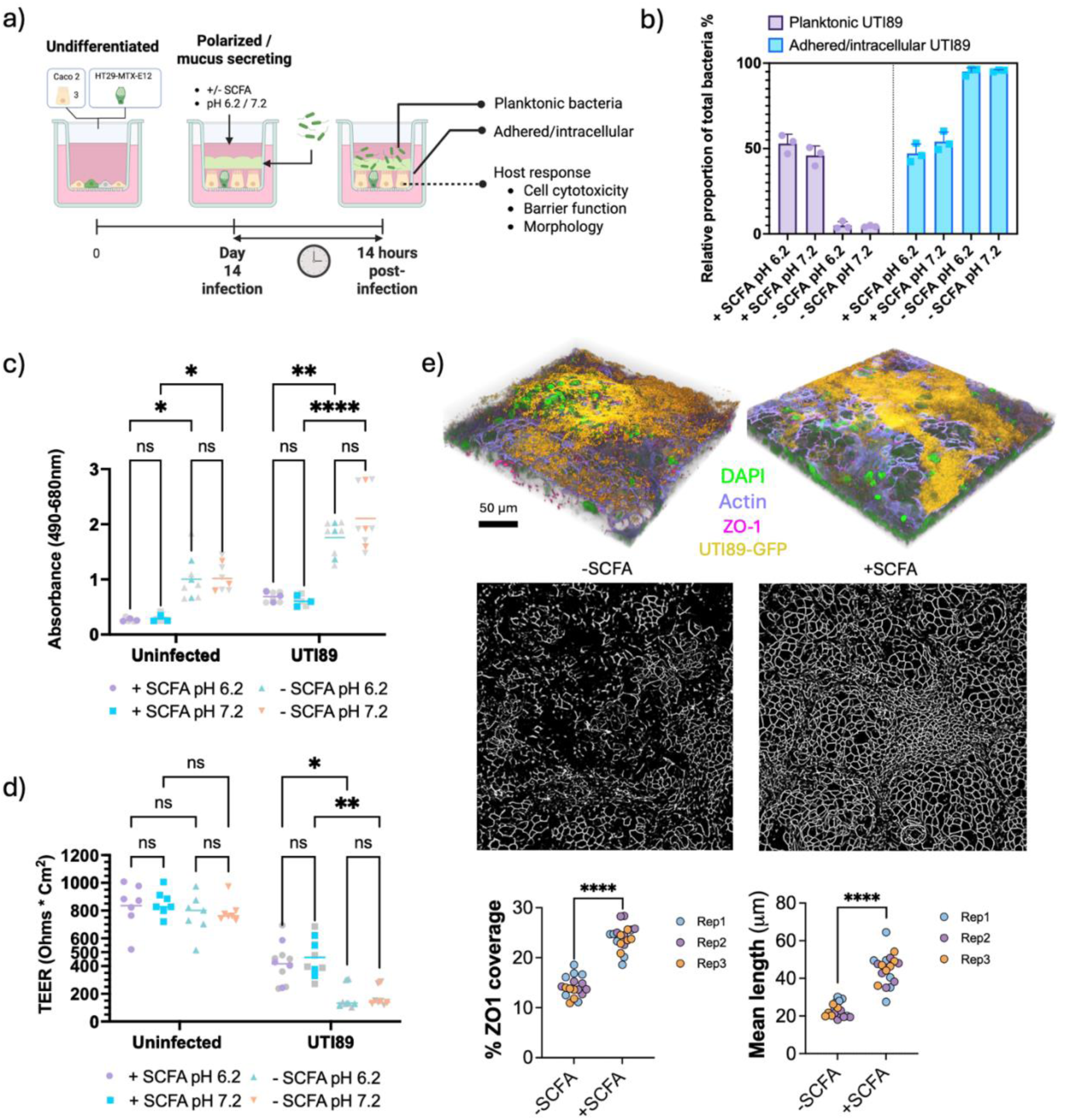
SCFAs treatment limits UPEC-induced epithelial barrier damage and modulates host-pathogen dynamics. **a)** Schematic of the experimental workflow. Created in BioRender. Yuen, C. (2026) https://BioRender.com/a2dqsta **b)** Relative proportion of bacterial burden in planktonic and epithelial-associated fractions following infection with UTI89 under SCFA-treated and untreated conditions at pH 6.2 and 7.2. Each point represents an independent biological replicate. **C)** Baseline epithelial cytotoxicity measured by LDH release following 14 h of infection. Data are presented as absorbance at 490/680 nm. Each point represents an independent biological replicate (coloured) averaged from 2 technical replicates (grey). Statistical significance was assessed by ordinary 2-way ANOVA: ns = not significant; *p<0.05; **p<0.01, ***p<0.001; ****p<0.0001. **d)** Transepithelial electrical resistance (TEER) measurements 14-hour post infection. Each point represents an independent biological replicate (coloured) with 2 technical replicates (grey). Statistical significance was assessed as in c). **e)** Representative 3D reconstructions (nuclei [green], F-actin [purple], ZO-1 [Magenta], and UTI89-GFP [yellow] are shown) and corresponding segmentation masks of ZO-1 tight junction staining in UTI89-infected monolayers at pH 6.2 in the absence or presence of SCFAs. Quantification of ZO-1 mask coverage (% area) demonstrates increased tight junction coverage in SCFA-treated monolayers. Points represent measurements from independent fields of view across three biological replicates (6 regions imaged per biological replicate). Statistical significance was assessed as in c).

To assess whether SCFAs modulate epithelial cytotoxicity in response to UPEC challenge, lactate dehydrogenase (LDH) release was quantified after 14 h of infection under SCFA-treated and untreated conditions at pH 6.2 and 7.2 (**Fig. 5C**). SCFA treatment alone significantly decreased LDH release in uninfected epithelial monolayers, indicating that SCFAs are not intrinsically cytotoxic, but indeed may have a baseline protective effect.

During UTI89 challenge, SCFA treatment resulted in an approximately two-fold decrease in LDH release independent of pH. These findings suggest that SCFAs prevent infection-associated cytotoxicity and may reduce the overall magnitude of epithelial damage incurred during UPEC challenge, supporting a role for SCFAs in enhancing epithelial tolerance to infection.

We further evaluated the impact of SCFA treatment on epithelial barrier integrity during UPEC challenge by measuring transepithelial electrical resistance (TEER) under SCFA-treated and untreated conditions at pH 6.2 and 7.2 (**Fig. 5D**) post-infection. In uninfected epithelial monolayers, SCFA treatment did not significantly alter TEER values, indicating that SCFAs alone do not affect baseline barrier integrity. Infection with UTI89 resulted in a reduction in TEER relative to uninfected controls, suggesting infection-induced barrier disruption. Across both pH conditions, SCFA-treated monolayers maintained higher TEER values following infection compared with untreated infected controls, suggesting preservation of barrier integrity via treatment with SCFAs. Consistent with our functional measurement, image analysis of confocal maximum-projections further revealed enhanced ZO-1 tight-junction coverage across our SCFA treated monolayers following infection, supporting improved maintenance of epithelial junction architecture during UPEC challenge (**Fig. 5E)**.

Together, these results demonstrate that, in the co-MIMIC model system, UPEC strain UTI89 induces epithelial damage characterized by compromised barrier integrity and increased cytotoxicity. The consistency of these findings across morphological, barrier (TEER), and cytotoxicity (LDH) readouts strengthens the conclusion that epithelial disruption is a direct consequence of UPEC infection under the tested conditions. Moreover, SCFA treatment significantly mitigated infection-associated epithelial damage, supporting a role for SCFAs in limiting UPEC-induced epithelial dysfunction.

## Discussion

Recurrent urinary tract infections (rUTI) remain a major clinical challenge, a problem compounded by the escalating AMR crisis (Von Vietinghoff et al., 2024). The gastrointestinal tract is increasingly recognised as a critical reservoir for uropathogens, probably enabling repeated transmission to the urinary tract (Schembri et al., 2022).

Mounting evidence supports a gut-bladder transmission model, in which pathogens shuttle from gut to bladder either via the bloodstream or, more commonly accepted, the faecal-periurethral-bladder route (Salazar et al., 2022; Schembri et al., 2022). Despite this, the mechanisms by which the intestinal microenvironment modulates uropathogen persistence/ virulence and, by proxy, UTI susceptibility, remain poorly understood.

Key gut microbial taxa responsible for fermenting dietary fibre into SCFAs are consistently found at reduced abundance in patients with rUTI, underscoring the potential role of both microbial composition and metabolite availability in rUTI pathophysiology (Worby et al., 2022b; H. J. Yang et al., 2022). While SCFAs are known to modulate virulence in *E. coli* and other enteric pathogens (Kadry et al., 2023; Lawhon et al., 2002; Zhang et al., 2020), their direct influence on UPEC colonisation traits within the gut niche has not been fully defined. In this study, we examined this question across key UPEC strains including clinical isolates, both from the urine of symptomatic and asymptomatic individuals. We show that SCFAs suppress critical UPEC colonisation behaviours, including swimming motility and type 1 pilus phase variation, and impose a pH-dependent delay in bacterial growth. By leveraging a polarised mucus secreting human intestinal model, co-MIMIC, we further demonstrate that SCFAs mediate protection of the gut barrier in response to UPEC colonisation. These findings offer mechanistic insight into how healthy gut-derived metabolites may influence UPEC colonisation and virulence in vitro, providing a foundation for understanding potential effects on intestinal colonization dynamics in the context of rUTI susceptibility.

### Acidified SCFAs significantly impact the lag phase of growth and may trigger metabolic adaptation

The lag phase of growth is a dynamic and adaptive period during which bacteria are introduced into a new environment and are considered non-replicative (Bertrand, 2019). In this study, we observed that colonic concentrations of SCFAs significantly prolonged the lag phase of UTI89 growth under acidic conditions. We propose that, similar to some other enteric pathogens, protonated SCFAs may affect bacterial cells more readily at lower pH, leading to intracellular acidification and activation of acid stress responses, as has been previously reported (Sorbara et al., 2019). Such stress likely triggers metabolic and transcriptional reprogramming before cells resume exponential growth (Bertrand, 2019). This adaptation period could be a critical window during which colonisation-related gene expression is altered in UPEC. Interestingly, commensal SCFAs have also been shown to disrupt lipid membrane homeostasis in *Staphylococcus aureus* with effects on bacterial growth, membrane integrity, attenuation of accessory genes and enhanced sensitivity to antimicrobials (Fletcher et al., 2025). Despite the effect on the lag phase, we observed no significant difference in the final carrying capacity. As such, final bacterial quantity at endpoint may not reflect dynamic changes during growth that nevertheless influence colonisation potential, particularly if there are transcriptomic shifts. Inconsistencies in bacterial abundance as a risk factor for UTI may stem from differences in microbial metabolic states. A clinical cohort study on rUTI patients (Young et al., 2024) found that *E. coli* strains in the rUTI gut exhibited a distinct transcriptional profile compared with those in non-rUTI individuals. Specifically, these bacteria shifted from fermentative metabolism toward aerobic metabolism. This shift suggests that *E. coli* in rUTI patients adapt to higher oxygen availability and oxidative stress in the gut environment. Increased levels of oxygen, nitrate, and reactive oxygen species (markers of intestinal inflammation) likely drive these transcriptional changes. Importantly, switching to aerobic metabolism may enable these bacteria to better survive and proliferate in a more oxygen-rich environment, including the periurethral area and the urinary tract, potentially increasing the risk of UTI recurrence.

Although bacterial intracellular acidification is one potential mechanism involved in responding to the SCFA-enriched or non-enriched environments, it has been previously demonstrated that protonated acetate can interact with two-component systems, such as the histidine kinase sensor BarA (Alvarez et al., 2021). The BarA/UvrY two-component system, with conserved homology amongst other bacterial genera such as Pseudomonas (*GacA*), Vibrio (*VarA*) and Salmonella (*SirA*), is a central regulatory element which controls transcription of CsrB/C non-coding RNAs, leading to sequestration of the global regulator CsrA. Given that CsrA positively regulates the flagellar master regulator *flhDC* and *fimA* transcription, SCFA-mediated activation of BarA/UvrY could represent a central signaling hub integrating metabolic state with motility and adhesion phenotypes. Loss of the UvrY response regulator reduces UPEC fitness in murine UTI models and decreases uroepithelial cell invasion (Palaniyandi et al., 2012), highlighting its role in infection. In *Salmonella*, the homologous *BarA*/*SirA* system modulates virulence genes in response to external SCFA signals, affecting invasion (Lawhon et al., 2002). These data suggest that *BarA*-*UvrY* may facilitate UPEC sensing of host metabolites like SCFAs to regulate virulence and adaptation during UTI, making it a potential drug target to disrupt infection. These mechanisms of action further highlight the potential role that SCFAs may play in facilitating metabolic rewiring and virulence regulation of enteric pathogens in the intestinal niche.

### The role of type 1 pili and motility in intestinal colonisation

While UPEC-mediated bladder colonisation is well characterised, the mechanisms underlying intestinal colonisation remain less well understood. Recent studies indicate that UPEC employ type 1, *Yeh*, and *Ucl* pili (F17-like pili) to facilitate intestinal colonisation in murine models and differentiated human epithelial cell lines (Azimzadeh et al., 2025; Spaulding et al., 2017). More recently, it has been shown that purified *YehD* and *UclD* lectin domains bind extensively to luminal and faecal content in a mouse model, whereas purified *FimH* binds more extensively throughout the secreted mucus layer and the bound mucosal layer (Azimzadeh et al., 2025). These findings highlight the role of FimH in mediating mucosal adhesion to enable UPEC to maintain residence within this secreted compartment. In the same study, deletion of *fimH* was further shown to result in a pro-flagellar phenotype mediated by *fliC* upregulation, potentially as a compensatory colonisation strategy. This upregulation may facilitate breaching of the inner mucus layer toward the epithelial surface, providing an advantage for persistence within the gut niche. FimH inhibitors have been shown to reduce both intestinal and bladder colonisation, highlighting them as a promising target for anti-adhesive therapies against UTI (Spaulding et al., 2017). One such development includes FimH-specific monoclonal antibodies, which were shown to protect against UPEC in murine models of urinary tract infection. This protection occurred through an Fc-independent mechanism in which the antibodies bind directly to the FimH adhesin protein with high affinity, recognizing conformation-dependent epitopes and thereby blocking FimH function (Lopatto et al., 2025).

To study the modulatory role of SCFAs on *fimH*, we leveraged a fim switch PCR assay. Regulation of type 1 pili is governed by phase variation, mediated by inversion of the fimS DNA element. The “ON” orientation of the *fimS* element is highly proportional to the transcription and expression of the downstream *fimA* and *fimH* region. Two site-specific recombinases, *FimB* and *FimE*, mediate this inversion process, and several global regulators, including H-NS, integration host factor (IHF), leucine-responsive regulatory protein (Lrp), LysR-type regulators, ppGpp, and OmpR, integrate environmental signals into this switch (Schwan, 2011). Therefore, in this study, we sought to determine if SCFAs could modulate this phase variable switch. Using the prototypical cystitis strain UTI89 and the pyelonephritis strain CFT073, we observed that acidified SCFAs significantly restricted inversion towards the *fimS* phase-ON orientation in a pH-dependent manner. This effect was observed under acidic conditions, consistent with effects observed in our growth kinetic analysis. Notably, our baseline (LB pH 7.2) phase-ON expression under phase-ON-inducing conditions was lower in CFT073 compared with UTI89, consistent with prior reports (Greene et al., 2015). We hypothesize the lower baseline “ON” levels in CFT073 may reflect its greater reliance on alternative adhesins, such as P pili.

Although type 1 fimbriae are an important factor in mediating UPEC colonization in the intestinal niche, flagellar mediated swimming motility may also play a role. In the urinary tract, motility can facilitate ascension toward the kidneys, while in the gut, *fliC*-mediated motility in uropathogenic *E. coli* has been implicated in facilitating breaching of the inner mucus layer towards the epithelium (Azimzadeh et al., 2025). Beyond locomotion, flagellin is a potent pathogen-associated molecular pattern (PAMP) recognised by Toll-like receptor 5 (TLR5) on intestinal epithelial cells and dendritic cells, triggering NF-κB–mediated pro-inflammatory cytokine production such as IL-8 (Steiner, 2007). This inflammatory response can disrupt epithelial integrity, recruit immune cells, and alter the gut environment in ways that may favour pathogen persistence.

In this study, we observed that SCFAs at colonic concentrations significantly attenuate swimming motility across a panel of UPEC strains, at neutral pH. In the gut, such suppression may not only limit bacterial penetration into the inner mucus layer and subsequent epithelial colonisation but also reduce exposure of host immune receptors to flagellin, thereby dampening inflammation and preserving mucosal barrier function. Taken together, although *fimH* and *fliC* regulation are considered inversely regulated, SCFAs may serve to repress both fimbriae and motility phenotypes.

### SCFAs limit UPEC-associated epithelial damage by modulating host-pathogen interactions

In this study, SCFA treatment favoured a planktonic bacterial lifestyle and was associated with reduced epithelial damage during UPEC infection. Following 14 h of infection, SCFA-supplemented conditions exhibited a relatively higher abundance of planktonic bacteria, whereas mucosal-associated bacteria were proportionally more prevalent in the absence of SCFAs. Consistent with this observation, infection with UPEC strain UTI89 significantly disrupted epithelial barrier integrity, as measured by transepithelial electrical resistance (TEER). In contrast, SCFA-treated monolayers maintained substantially higher TEER values compared with untreated controls, indicating preservation of barrier function during infection.

To contextualize these findings with our earlier observations that SCFAs influence bacterial lag-phase dynamics, we propose that during early stages of host colonization, UPEC likely experiences similar growth kinetics across conditions. Within this metabolic adaptation window, SCFA exposure may attenuate the expression or function of infection-associated virulence phenotypes, thereby limiting epithelial damage without substantially altering bacterial burden. The observed shift toward a planktonic bacterial state in the presence of SCFAs may further reduce intimate epithelial interactions that contribute to barrier disruption and cytotoxicity.

Importantly, the protective effects of SCFAs are unlikely to be exclusively bacterium driven. Beyond their impact on bacterial physiology, SCFAs, particularly butyrate, have been shown to directly enhance epithelial barrier integrity through mechanisms including AMP-activated protein kinase (AMPK) activation and histone deacetylase (HDAC) inhibition, resulting in improved tight junction assembly. Accordingly, the preservation of TEER and enhanced ZO-1 coverage observed in SCFA-treated monolayers likely reflect a combination of reduced bacterial insult and direct host epithelial fortification during infection.

Together, these findings highlight the protective influence of SCFAs on epithelial integrity during UPEC challenge and underscore the broader relevance of gut barrier maintenance in the context of recurrent urinary tract infections (rUTIs). Increasing evidence suggests that gut-resident uropathogens may influence bladder pathology indirectly through systemic dissemination of bacterial products, including endotoxins and other pro-inflammatory mediators. Such circulating factors may sensitize distal mucosal sites, compromise urothelial defenses, and promote an inflammatory environment conducive to recurrent cystitis and lower urinary tract dysfunction. In support of this concept, women with rUTI have been reported to exhibit a distinct immune profile, including elevated levels of eotaxin-1, a gut-associated inflammatory chemokine also implicated in inflammatory bowel disease and Crohn’s disease (Worby et al., 2022b). Collectively, these observations suggest that SCFA-mediated preservation of gut barrier integrity may have implications beyond the intestine, potentially influencing susceptibility to recurrent urinary tract infection.

## Conclusion: A two-pronged mechanism

We hypothesize a two-pronged mechanism for SCFA-mediated protection, which may modulate both bacterial pathoadaptive behaviours whilst also conferring host protection (**Figure 6**). Our findings demonstrate that physiologically relevant concentrations of short-chain fatty acids (SCFAs) modulate key colonisation traits of uropathogenic *Escherichia coli* (UPEC), including *fliC* mediated swimming motility and *fimS* orientation directly. By leveraging the Co-MIMIC model, we further observe SCFA mediated modulation of UPEC host-pathogen dynamics at the intestinal epithelial interface. These effects, consistent with previous observations of SCFAs shaping the pathogenic potential of enteric bacteria, suggest that SCFA-mediated signalling within the gut could influence UPEC persistence and potentially the risk of rUTI.

**Figure 6.**
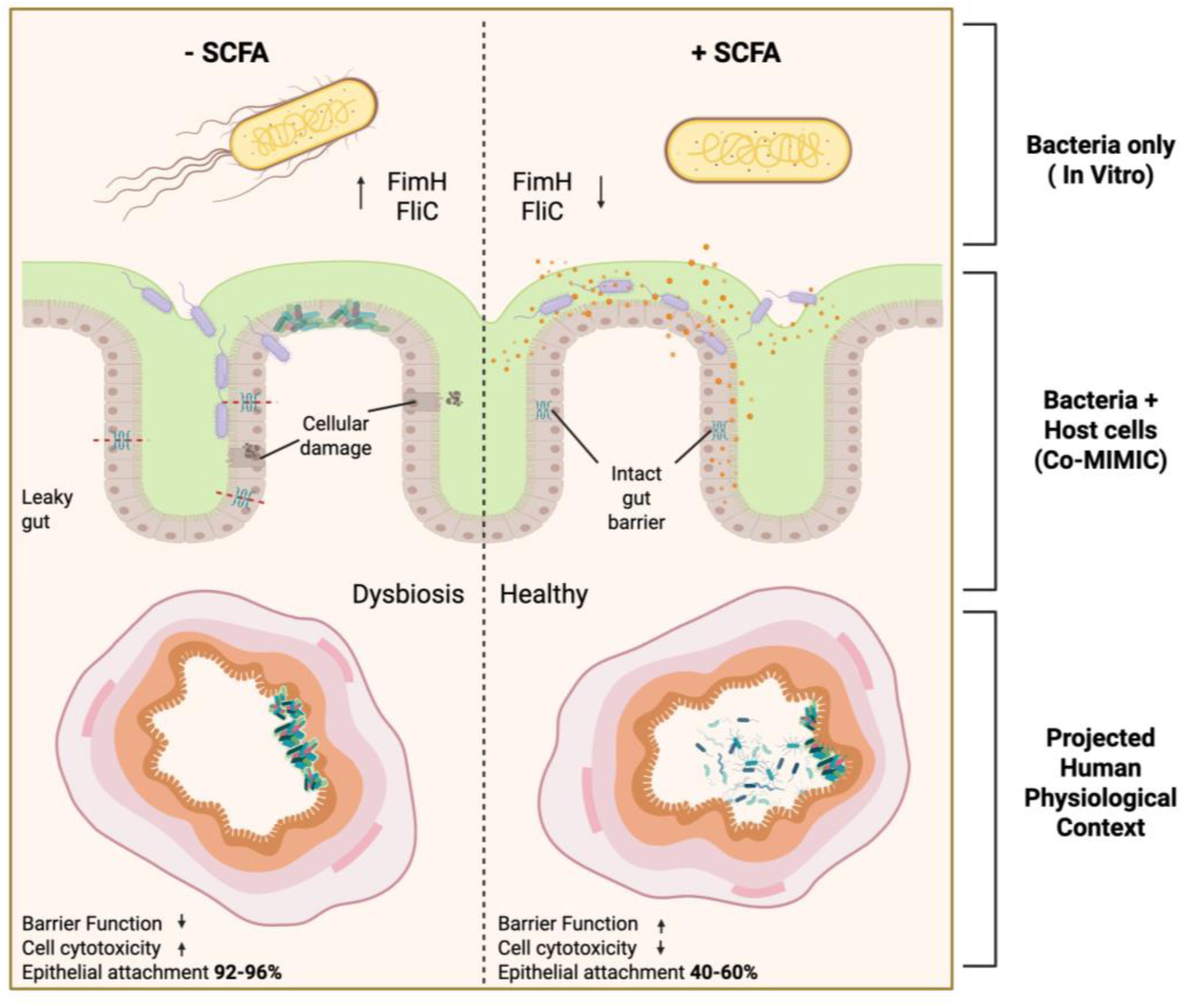
Conceptual model of SCFA-associated modulation of UPEC–host interactions. The schematic summarizes findings from bacteria-only, epithelial cells (Co-MIMIC) challenged with UTI89, and translationally projected contexts, depicting how SCFA availability is associated with altered UPEC colonization traits and improved epithelial barrier integrity. Created in BioRender. Yuen, C. (2026) https://BioRender.com/69af0ya

While this study provides mechanistic insight into how SCFAs modulate UPEC colonization in the intestinal niche, several limitations should be acknowledged. The polarized mucus-secreting epithelial model used here captures key features of the intestinal interface but lacks *in vivo* mucus thickness, immune cells, and complex polymicrobial communities that shape both SCFA metabolism and pathogen behavior *in vivo*. Moreover, the experimental SCFA mixtures and pH conditions simplify the complex spatial and dynamic temporal gradients present in the colonic lumen. We also note no differences in mucosal attachment in response to SCFA treatment and hypothesize that this may be attributed to bacterial saturation due to a long infection period as well as an absence of flow or natural turnover of the secreted mucosa (peristalsis). Additionally, the present work is limited to investigating SCFA-mediated effects on UPEC within the intestinal niche and does not address potential distal consequences in the bladder. Future studies employing defined microbial consortia or gnotobiotic models will be essential to determine how microbiota composition, dietary fiber availability, and SCFA flux influence UPEC adaptation and persistence within the gut reservoir. At the molecular level, identifying the specific signaling pathways through which SCFAs modulate host-pathogen interactions will refine our understanding.

In conclusion, given that gut colonization likely serves as a critical reservoir for rUTI, our use of an *in vitro* intestinal infection model provides mechanistic insights into how SCFAs have the potential to modulate UPEC colonization at the intestinal interface. In addition, this may have translational potential in informing how dietary patterns, microbiota composition, or therapeutic modulation of SCFAs might be leveraged to reduce UPEC intestinal carriage. Therapeutic approaches aimed at enhancing colonic SCFA production, such as increased dietary fiber intake, targeted prebiotics, or microbiota restoration may offer promising adjunct strategies to decrease UPEC persistence and virulence in the gut and mitigate rUTI recurrence. Future *in vivo* and clinical studies are needed to evaluate whether targeted manipulation of SCFA profiles can serve as an effective preventive strategy against UPEC-related infections.

## Acknowledgments

We would like to express our sincere gratitude to the Scott Hultgren lab for providing the WT UTI89 and left inverted repeat mutant, “Locked ON” UTI89 strain. We also thank Dr Davide Di Grande for 3D printing a modified platform for stable TEER analysis. We thank Dr Luke Allsopp and Dr Meriem El Keroui for critical review of the manuscript. We also thank Prof Meera Unnikrishnan for helpful discussion. Finally, we acknowledge funding from the Urology Foundation: innovation and research award.

