## Supplementary Data for "Gut-relevant short-chain fatty acids modulate host-pathogen dynamics of uropathogenic *Escherichia coli* at the colonic epithelial interface"

### Supplementary Figure Legends

**Supplementary figure S1 | TrackMate pipeline for estimation of extrapolated nuclei count across the 1.12 cm$^2$ polycarbonate membrane. a)** individual nuclei imaged via confocal microscopy at 40x magnification (blue). b) Circles (magenta) indicate identification of individual nuclei via LAP tracker. c) Converted tracks counted as determined by Trackmate plugin. Each coloured line represents a single tracked nucleus. d) Number of tracks were quantified from 4 different regions of interest across 3 independent experiments and extrapolated to estimate cell density across a 1.12 cm$^2$ polycarbonate membrane.

**Supplementary figure S2 | SCFA mixture and individual SCFAS (Acetate, Butyrate and Propionate) modulate growth kinetics of uropathogenic *E. coli* strain UTI89 in a pH- and concentration-dependent manner.** Growth kinetic analysis of Uropathogenic *E. coli* strain UTI89 grown in SCFA mixture (Acetate: Butyrate: Propionate, 60:20:20 molar ratio), Acetate (120 mM, 60 mM, 20 mM), Butyrate (120 mM, 60 mM, 20 mM) and Propionate (120 mM, 60 mM, 20 mM) at both pH 6.2 (**a**) and pH 7.2 (**b**). Error bars shown as ± standard deviation of the mean.

**Supplementary figure S3 | Quantitative (non-parametric fitting) comparison of bacterial growth parameters under different pH and short-chain fatty acid (SCFA) conditions**. The following parameters are displayed in log10 relative units: maximum specific growth rate (μ_max, blue), doubling time (t_D, purple), lag phase duration (λ, pink), maximum optical density (y_max, red), change in optical density (ΔY, orange), time to maximum growth rate (t(μ_max)) and area under the curve (dark blue) for UTI89 cultures grown in NaCl or SCFAsupplemented medium at neutral (7.2) and mildly acidic (6.2) pH. Statistical significance is indicated above the bars: ns = not significant; *p<0.05; **p<0.01, ***p<0.001; ****p<0.0001. (c) Table of mean ± standard deviation for each growth parameter in UTI89 under the indicated conditions. Data include μ_max ($h^{-1}$), t_D (h), λ (h), y_max ($OD_{600}$), and ΔY ($OD_{600}$) for NaCl and SCFA mixture (120 mM) treatments at pH 7.2 and 6.2. All data shown here represent 3 biological and 3 technical replicates.

**Supplementary figure S4 | Example of Parametric and non-parametric modelling of UPEC growth under SCFA conditions at pH 7.2 and 6.2.** Growth in the presence of both SCFAs and NaCl was fitted using the appropriate parametric model. a) The top panels display observed bacterial growth over time with the corresponding model fits (green curves), showing differences in the shape and saturation of the growth trajectories. b) The bottom panels represent non-parametric analyses, where smoothed estimates of growth rates (solid blue lines) are plotted against the fitted rate curves (dashed blue lines). (Graphs are representative of 3 biological replicates and 3 technical replicates)

**Supplementary figure S5 | fimS PCR assay to determine %ON/%OFF amplicons in UTI89 and CFT073 under SCFA supplemented conditions (pH 6.2/7.2).** a) Schematic outlining methodological workflow. Strains were grown in their respective media (+/- SCFA at pH 6.2 and 7.2) in phase-ON induced conditions. Phase-ON induction was achieved by static incubation at 37 °C for 24 hours, followed by a 1:1000 subculture and a subsequent 24-hour static incubation at 37 °C, as previously described (Greene et al., 2015). From this, we normalized our bacterial samples to $OD_{600}$ = 1 and 1 ul was used as DNA input for fimS PCR amplification. Restriction digestion followed by gel electrophoresis was used to separate individual fragments and imageJ analysis was used to measure fluorescence intensity. b) Schematic describing fimS amplicon in the "ON" (160 bp and 442 bp) and "OFF" (404 bp and 198 bp) orientation. Agarose gels of separated fragments in all conditions for c) UTI89 and d) CFT073. Experiments represent n=5 biological replicates.

**Supplementary figure S6 |** Cell Tracker green staining of HT29-MTX-E12 goblet-like cells indicating formation of cell clustering and "island" structures surrounded by Caco2 enterocytes. DAPI (Cyan), F-Actin (Magenta), HT29-MTX-E12 (Green) b) Mucus characterization of Co-MIMIC model indicating MUC2 expression colocalized with HT29-MTX-E12 goblet cells. Acidic mucins on parallel cultures confirmed via alcian blue staining (Right). DAPI (Cyan), MUC2 (Red).

**Supplementary figure S7 | CFU/ml bacterial quantification 14 hpi with UTI89 with or without SCFA treatment at pH 6.2/7.2.** a) Planktonic as well as Adhered/ intracellular (Epithelial) fractions were quantified to determine bacterial localisation in response to SCFA treatment. Data represent 3 biological replicates.

**Supplementary Figures**

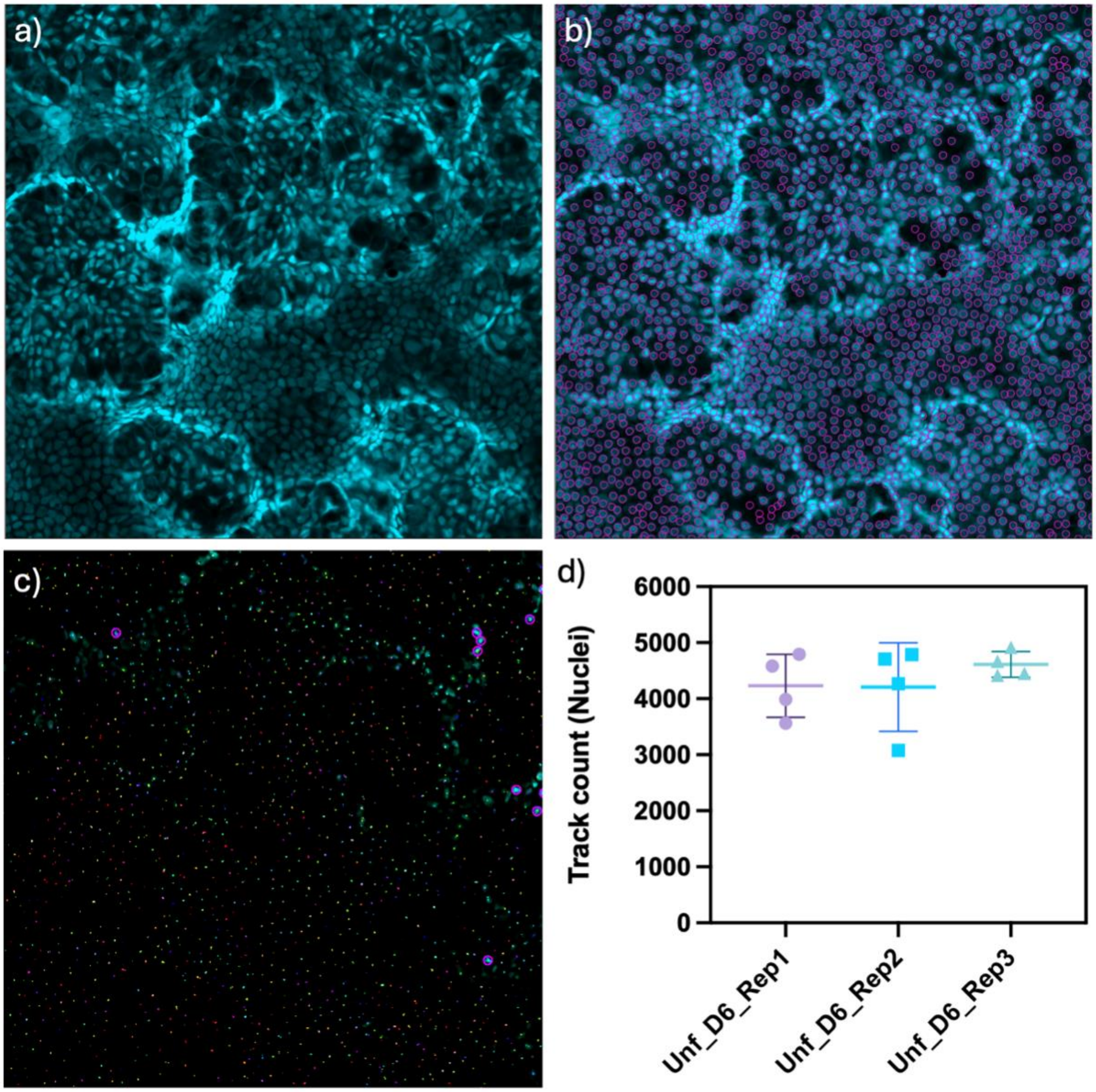


**Supplementary figure S1**

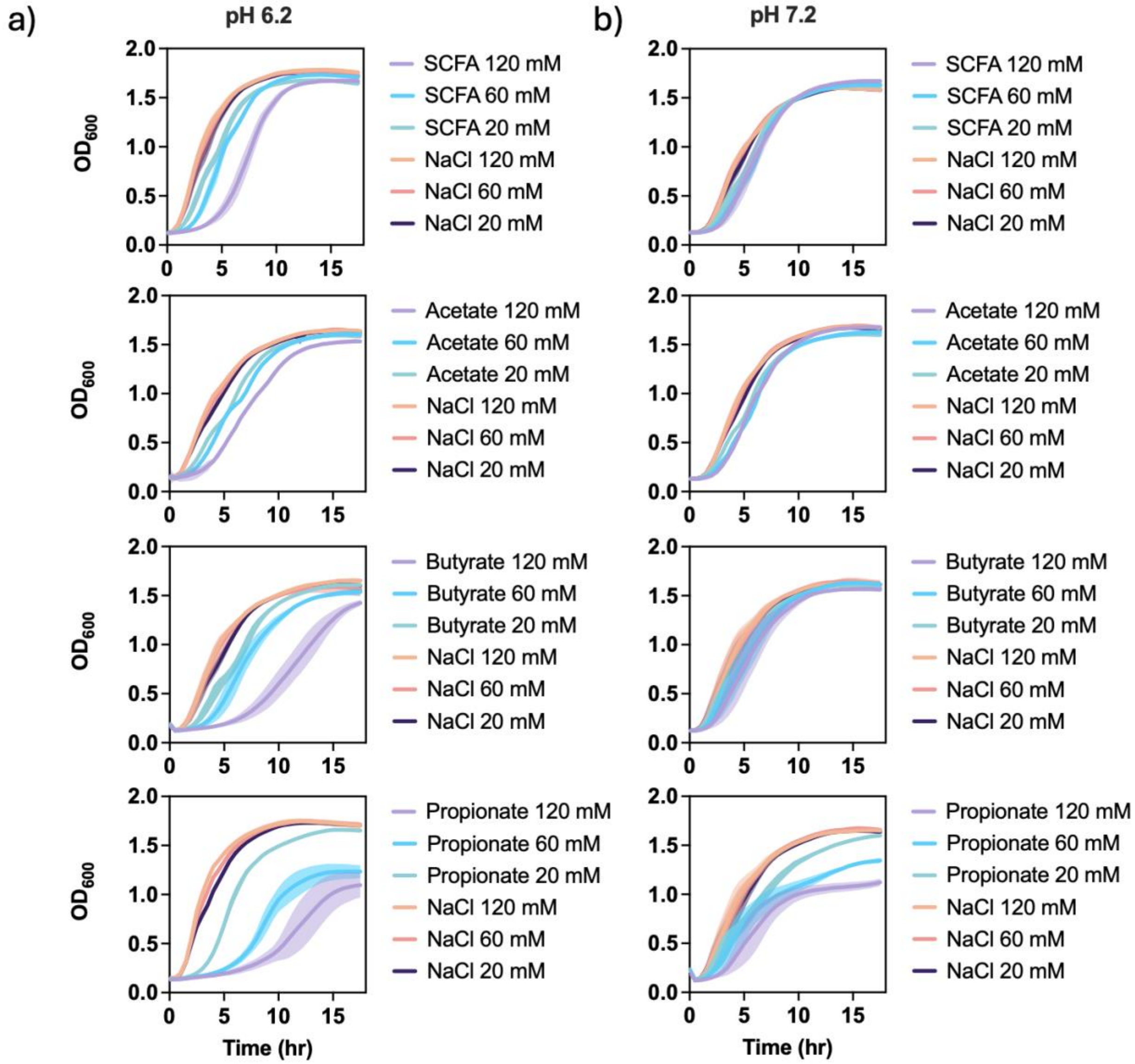


**Supplementary figure S2**

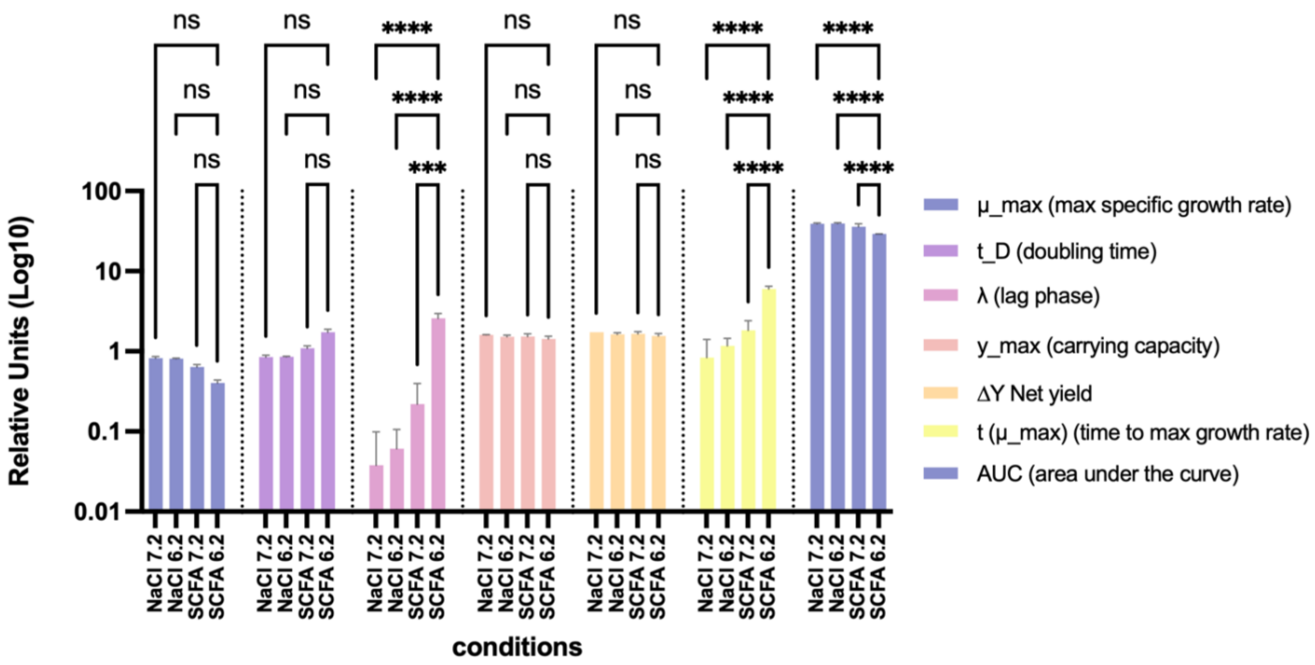


**Supplementary figure S3**

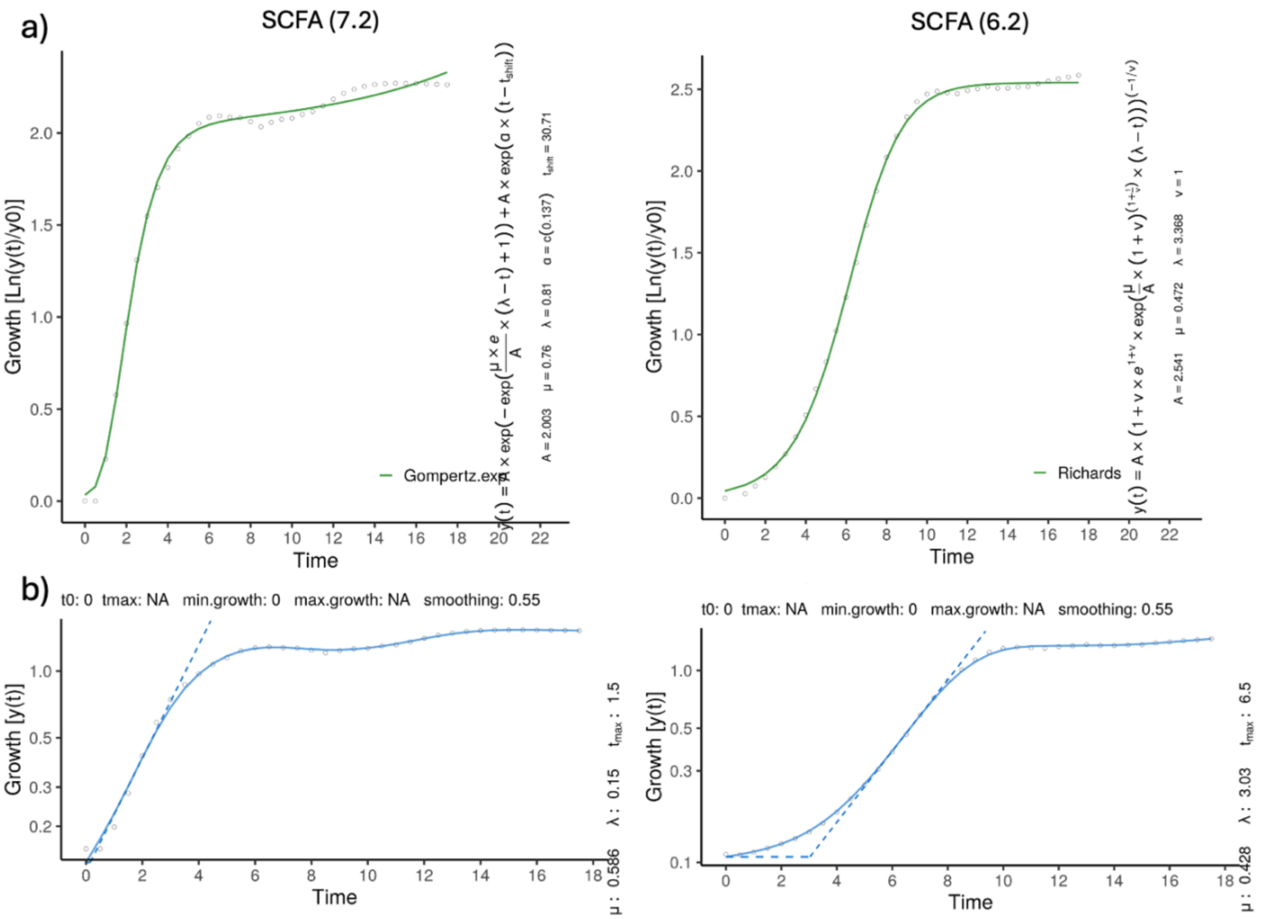


**Supplementary figure S4**

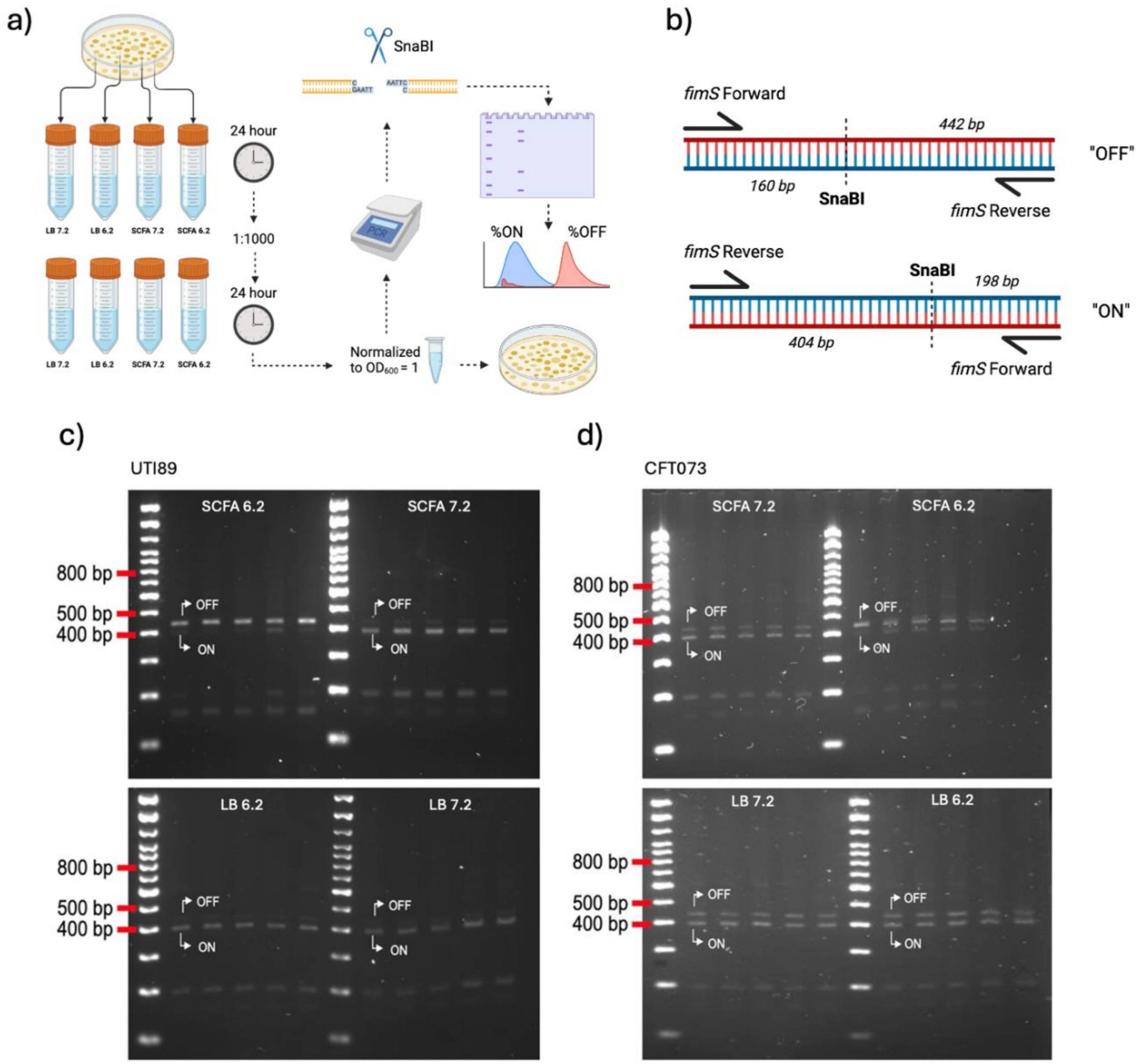


**Supplementary figure S5**

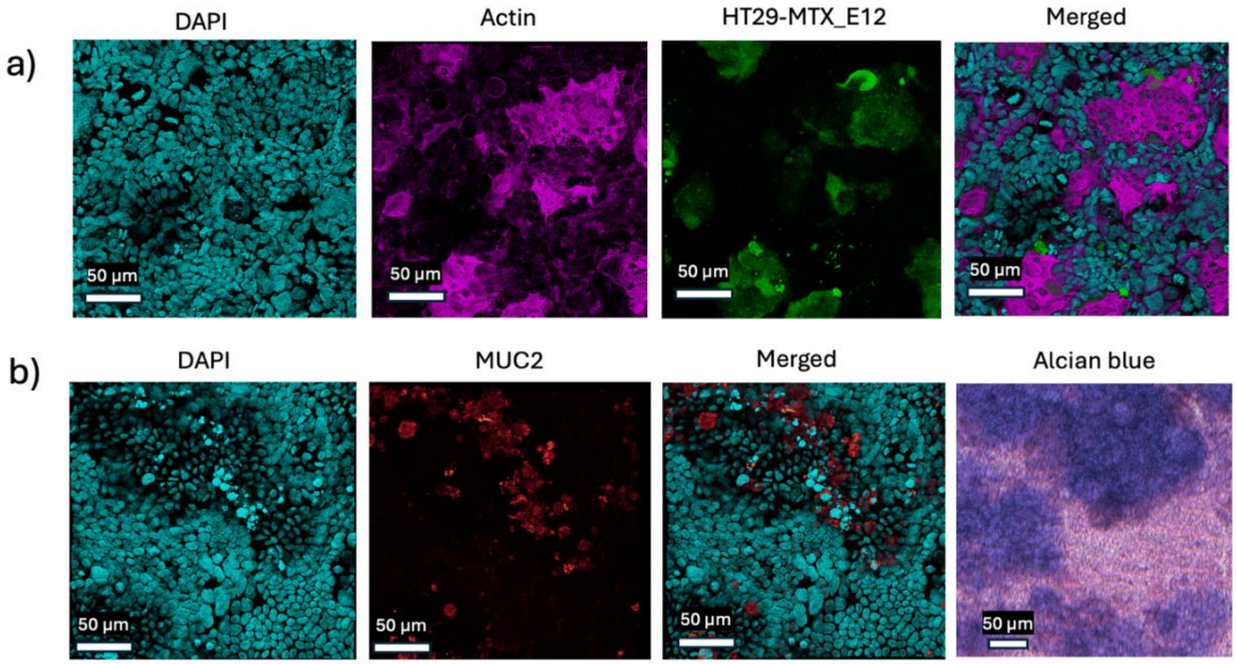


**Supplementary figure S6**

a)

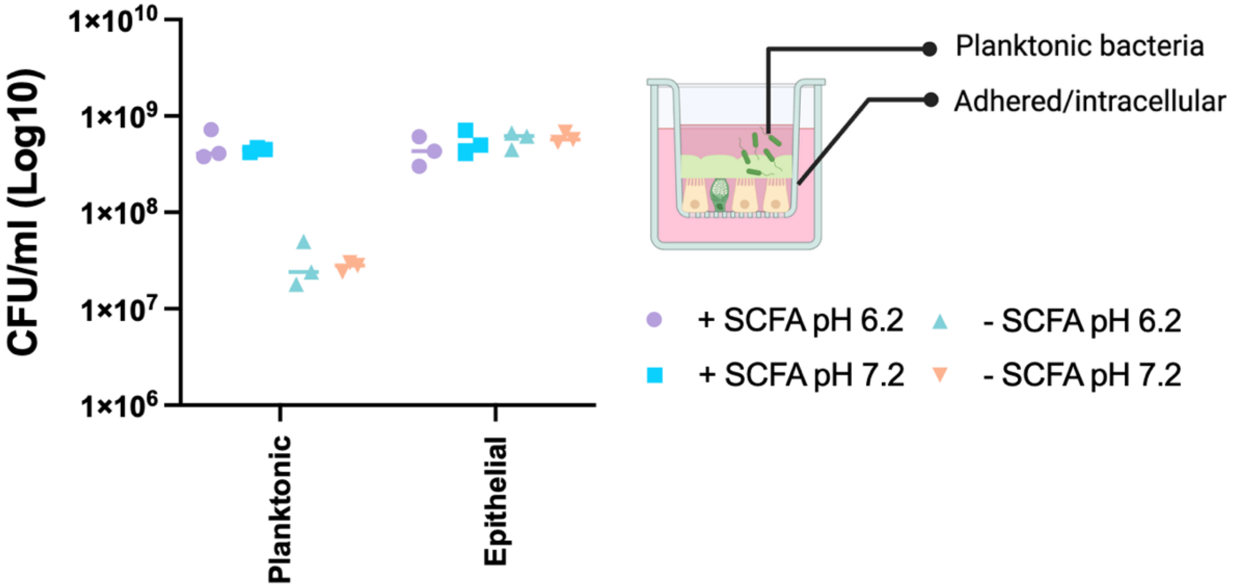


**Supplementary figure S7**
